# Divergent Functions of Late ESCRT Components in *Giardia lamblia*: Insights from Subcellular Distributions and Protein Interactions

**DOI:** 10.1101/2024.10.25.620214

**Authors:** Nabanita Patra, Nabanita Saha, Trisha Ghosh, Pritha Mandal, Avishikta Chatterjee, Babai Hazra, Sandipan Ganguly, Srimonti Sarkar

## Abstract

*Giardia lamblia*, a human gut pathogen, possesses a minimal ESCRT (Endosomal Sorting Complex Required for Transport) machinery. Paradoxically, there are multiple paralogs of some late-ESCRT components. There are three paralogs for Vps4, GlVps4a, GlVps4b, and GlVps4c, and two for Vps46, GlVps46a, and GlVps46b. This study addressed whether these paralogs discharge overlapping and/or distinct cellular functions by determining the sub-cellular distribution of the paralogs in trophozoites and during encysting. Consistent with the distribution of orthologs in model organisms, most of these components were found to be associated with various cellular membranes, particularly in regions of acute membrane bending. Some of these paralogs are also associated with microtubule structures, such as cytoplasmic axonemes and the median body. Considering their diverse sub-cellular distributions, it is likely that they perform non-overlapping functions within the cell. Furthermore, their redistribution during encystation indicates that they may play a role in the morphological and functional changes accompanying this transition. The study also characterized GlIst1, an ESCRT-III accessory protein that undergoes unique post-translational myristoylation at lysine 43, potentially aiding its membrane recruitment. GlIst1 selectively interacts with GlVps4b through non-canonical MIT-MIM interactions. GlIst1 also exhibits selective interaction with GlVps46b. Such selective interaction of GlIst1 with only specific paralogs of GlVps4 and GlVps46 further underscores the distinct cellular roles of these paralogs.

**Author Summary:** *Giardia lamblia*, a unicellular protozoan parasite, is the causative agent of giardiasis, a water-transmitted disease affecting millions globally. This disease poses a substantial threat to public health, especially in less developed countries, where clean water and proper sanitation are scarce. The parasite manifests in two morphologically distinct forms, trophozoites and cysts. Transformation between these forms is essential for the organism’s survival, spread, and infection processes. Trophozoites, the active and motile form of *Giardia*, inhabit the small intestine of the host and trigger infections. These trophozoites can transform into cysts through encystation, enabling the parasite to endure harsh external environments and spread between hosts through contaminated water or food sources. The transition between these states necessitates extensive membrane restructuring. Such changes are likely to involve the Endosomal Sorting Complex Required for Transport (ESCRT) machinery, as it has been shown to participate in both prokaryotic and eukaryotic membrane remodeling events. Our research sheds light on the ESCRT machinery in *G. lamblia*, a crucial membrane-shaping system that may facilitate the transition between trophozoites and cysts. The ESCRT machinery in *G. lamblia* is distinct from that in yeast and humans, representing one of the most basic ESCRT systems. Our investigation provides valuable information about the intracellular distribution of various late-ESCRT components under different conditions, their potential functions, and their interactions with other late-ESCRT components. These findings may contribute significantly to our understanding of the basic operation of the ESCRT machinery in this parasite.

## Introduction

Membrane deformation is an integral part of several cellular processes, and the ESCRT machinery is one of the prime architects of several membrane remodeling events [1]. It plays a pivotal role in diverse cellular processes, including multivesicular body (MVB) formation, tubular endosomal trafficking for the recycling of receptors, cytokinesis, viral budding, plasma membrane repair etc. [2, 3]. It was first discovered in *Saccharomyces cerevisiae*, where a screen for vacuolar protein sorting mutants identified many of the constituents of this machinery [4]. Subsequent studies have shown that ESCRT-mediated membrane remodeling is evolutionarily conserved as ESCRT-related components are present in archaea and bacteria [5, 6]. This indicates that the origin of the ESCRT machinery predates the emergence of eukaryotes.

The Membrane recruitment of ESCRT components is sequential, and the precise orchestration of this process ensures efficient cargo sorting and membrane remodeling [7]. In the case of endosomal sorting, while the early ESCRT complexes (ESCRT-0, -I and -II) are involved in the recognition and subsequent aggregation of cargo proteins, the ESCRT-III complex, which functions downstream of the first three, plays a pivotal role in the final membrane remodeling steps [8]. For this pupose, the ESCRT-III components polymerize into spiral filaments on the membrane surface [9, 10]. Sequential recruitment and removal of various ESCRT-III components causes a change in the shape of this spiral which facilitates membrane invagination and scission [11, 12]. This membrane deformation and subunit exchange is fueled by ATP hydrolysis by the AAA-ATPase Vps4 and its associated proteins that form the Vps4 complex [13]. This stepwise assembly and disassembly of ESCRT components on the membrane ensures the precise spatial and temporal control of membrane remodeling events [14, 15].

It is important to note that not all ESCRT-dependent cellular processes require participation of all ESCRT components. For example, ubiquitinated cargo recognition on the endosomal surface and the subsequent sorting into the vacuole lumen requires ESCRT-0 and –I [16, 17]. However, plasma membrane repair, cytokinetic abscission, virus and microvesicle budding from the plasma membrane occur without the involvement of ESCRT-0 and ESCRT-I. Instead, they are dependent on the ESCRT-III and Vps4 complexes, which together are sufficient for constricting and severing the narrow membrane neck [18]. Furthermore, certain membrane deformation events can occur with some, but not all, of the ESCRT-III components. For example, Ist1 and CHMP1A/CHMP1B are the main ESCRT-III family members driving cytokinesis [19], while CHMP4 (Snf7) and CHMP2 (Vps2) filamentous structures drive membrane scission during HIV budding from mammalian cells [20, 21], and the repair of tears in the plasma membrane involves CHMP2A, CHMP3, CHMP4B along with Vps4. Membrane reshaping by ESCRT-III components can occur even without Vps4, as is the case for sorting of the mannose 6-phosphate and transferrin receptors through the formation of endosomal tubules [2]. The formation of these endosomal tubules requires the late-ESCRT components, Ist1 and CHMP1B, which function in conjunction with spastin, to efficiently divide endosomal tubules [2]. Spastin is similar to Vps4 as it contains both a AAA ATPase and an Microtubule Interacting and Trafficking (MIT) domain; it uses the latter to interact with the aforementioned ESCRT components [22, 23]. However, unlike Vps4, it has the ability to severe microtubules [24]. Thus, the ESCRT machinery driving membrane sculpting is highly modular, wherein only some components can function at a certain cellular location to bring about the desired outcome.

The complexity of the ESCRT machinery has evolved with increasing complexity of organisms. In general, complex life forms have ESCRT complexes that are composed of more components, including many that are paralogous, whereas simpler organisms have less elaborate ESCRT machinery with fewer components. One of the simplest eukaryotic ESCRT machinery is that of *Giardia lamblia* (also termed *G. intestinalis* or *G. duodenalis*), a human gut pathogen [25]. This parasite has two morphologically distinct forms: trophozoites and cysts [26, 27]. The transition between them is crucial for its survival within the host and transmission from one host to another [28]. This transformation necessitates extensive membrane restructuring [29, 30], and ESCRTs are likely to be involved in such processes. Previous research indicates that *G. lamblia* possesses a minimal ESCRT machinery, lacking the early ESCRT complexes, ESCRT-0 and ESCRT-I [31]. In addition, the ESCRT-III complex of *G. lamblia* comprises fewer core and accessory components, including Vps20, Vps2, and Vps24 as core ESCRT-III components, and Ist1 and the paralogs of Vps46 as accessory components. The ESCRT-III subunits Snf7, Vps60, and Vta1 are likely to be absent in this parasite. However, some late-ESCRT components are present in multiple copies, notably Vps4 (GlVps4a, GlVps4b, and GlVps4c) and Vps46 (GlVps46a and Vps46b) [25]. As mentioned previously, ESCRT-dependent membrane deformation can proceed without the entire set of ESCRT proteins. Hence, the absence of certain ESCRT components in *Giardia* is unlikely to render the ESCRT-dependent process non-functional in this parasite.

The presence of multiple paralogs of GlVps4 and GlVps46 suggests that these paralogs may discharge overlapping and/or distinct functions. To determine if these different paralogs of Vsp4 and Vps46 are functionally distinct, this study has immunolocalized these in trophozoites and encysting trophozoites. The results showed that these proteins localize to various membrane-deforming sites and microtubule-rich regions. They mostly occupy unique sites within the cell, except for a few regions, such as the peripheral vesicles (PVs), ventral disc (VD), flanges and flagellar axonemes, where more than one of these components are present. In addition, we report the characterization of the Ist1 ortholog in *Giardia* (GlIst1) and show that it undergoes myristoylation at an internal K residue that is unique to the *Giardia* protein. This post-translational modification is likely to aid in its membrane recruitment. Furthermore, since GlIst1 selectively interacts with only one Vps4 and one Vps46 paralog, myristoylation of GlIst1 may represent a new mechanism for the selective membrane recruitment of late-ESCRT components.

## Methods

### Axenic culture of *G. lamblia*

The axenic culture of *G. lamblia* (ATCC 50803/WB clone C6) trophozoites in TY-I-S-33 medium was carried out as previously described [32], and encystation was induced according to the protocol of [33].

### Construction of plasmids

For the yeast two-hybrid assay, the following ORFs were PCR amplified, using gene-specific primers listed in S1 Table, from *G. lamblia* genomic DNA: GL50803_101906 (*glvps4a*), Gl50803_16795 (*glvps4b*), Gl50803_15469 (*glvps4c*), Gl50803_15472 (*glvps46a*), Gl50803_24947 (*glvps46b*) and Gl50803_0011129 (*glist1*). The PCR products were cloned into the vectors pGAD424 and/or pGBT9 (Takara). The Primers used for cloning various fragments of *glist1* and *glvps4b* in these vectors are listed in S1 Table. The yeast genes, *VPS4* and *IST1* were PCR amplified from *S. cerevisiae* genomic DNA using a specific set of primer pairs (S1 Table) and cloned into pGAD424 and pGBT9, respectively. To generate antibodies for immunolocalization, *glvps4b*, *glvps4c*, *glvps46a* and *glvps46b* were PCR amplified using genomic DNA of *G. lamblia* (primers listed in S1 Table) and cloned into the expression vector, which was used for the expression of His-tagged protein. The constructs used for live-cell imaging were created by PCR amplification of *glist1* and yeast *IST1* from the respective genomic DNA with the corresponding primers (S1 Table) and cloned into pGOGFP426 and pRS425-RFP, respectively. All clones were sequenced to confirm the presence of inserts. The Details of all the constructs are given in S2 Table.

### Expression of proteins and generation of antibodies

His-tagged GlVps4b was expressed in *E. coli* by induction with 0.5 mM IPTG at 30 °C for 4 h. Following induction, cell lysis was carried out by sonication (20s pulses followed by 1 min gap at 75% amplitude). The extract was analyzed by SDS-PAGE to ensure the induction of the desired protein, and the His-tagged protein was purified using Ni-NTA (Qiagen). Similarly, His-tagged versions GlVps4c_1-261_, GlVps46a, and GlVps46b were expressed in *E. coli* by induction with 0.5 mM IPTG for 16 h at 20°C, 0.2 mM of IPTG for 3 h at 37°C, and 0.5 mM IPTG for 16 h at 20°C, respectively. The 6X-His-tagged proteins were purified as described previously and the purified proteins were handed over to BioBharti Lifesciences (Kolkata, India) for generation of the respective antibodies in rabbits.

### Protein extraction and western blotting

(A) *G. lamblia* extract was prepared by resuspending it in lysis buffer (50 mM Tris-Cl, 100 mM NaCl, 2% SDS, 1% Triton X-100, pH 8.0), followed by incubation on ice for 30 min. The protein fraction was then obtained by centrifugation at 12,000 rpm for 10 min, and the resulting supernatant was subjected to Bradford assay for quantification of extracted protein. For western blotting a 1:5000 dilution was used for antibodies against GlVps4a, GlVps4b, GlVps4c, and GlVps46b, whereas a 1:8000 dilution was used for the antibody against GlVps46a. All dilutions were prepared in 1X TBS containing 0.2% BSA. After incubating the membranes with the respective primary antibodies for 2 h, they were washed three times with 1X TBST and twice with 1X TBS. Following the washing, alkaline phosphatase-conjugated anti-rabbit antibody was used at a dilution of 1:5000 for GlVps4a, GlVps4b, and GlVps4c, whereas anti-rabbit antibody conjugated to AP was used at a dilution of 1:10000 for GlVps46a and 1:8000 for GlVps46b. After incubation with secondary antibodies for 2 h, the membranes were washed three times with 1X TBST for 5 min each, followed by two washes with 1X TBS. The blots were developed using NBT/BCIP. Similarly, total protein from yeast harbouring BD-GlVps46a, BD-GlVps46b was prepared by resuspending the transformants in suspension buffer (1 M Tris-Cl pH 7.5, 0.5 M EDTA, 2.5 M NaCl, NP-40, 1 M DTT, of 0.1 M PMSF and protease inhibitor cocktail) and vortexing in the presence of glass beads for 10 min at 4°C. Centrifugation was carried out at 13,000 rpm for 15 min, and the resultant supernatant was collected. The Bradford assay was used to quantify the extracted proteins. For western blotting, the membrane was blocked overnight with 2.5% BSA in 1X TBS. Antibodies against GlVps46a (1:8000 dilution) and GlVps46b (1:5000 dilution) was used. As loading control, anti-3-PGK (Invitrogen) was used at a 1:8000 dilution. The membranes were incubated with the respective antibodies for 2 h. Membranes were washed thrice with 1X TBST and twice with 1X TBS. Anti-mouse or anti-rabbit AP-conjugated secondary antibody (Abcam) was used at a 1:8000 dilution in 1X TBS for 1 h. Membranes were washed as previously described and developed using NBT/BCIP (Thermo Scientific).

### Immunolocalization

*Giardia* cells were harvested by chilling the tubes on ice for 5 min, followed by centrifugation at 1000g for 10 min. Cells were washed twice with 1X PBS and fixed with 4% formaldehyde for 15 min. Next, the cells were collected by centrifugation and treated with 0.8 M glycine in 1X PBS for 5 min at room temperature. The Cells were harvested by centrifugation and permeabilized with 0.2% Triton X-100 solution for 8 min. Next, the cells were blocked with 2% BSA for 1.5 h, harvested by centrifugation, and then incubated at 4°C overnight with primary antibody solution for GlVps4a (1:5000), GlVps4b (1:5000), GlVps4c (1:5000), GlVps46a (1:8000) or GlVps46b (1:5000) in 1X PBS. Cells were collected by centrifugation and washed thrice with 1X PBS, followed by incubation with a secondary antibody (anti-rabbit Alexa-488, 1:8000 dilution in 1X PBS) for 2 h. Then the cells were washed thrice with 1X PBS and resuspended in mounting medium containing p-phenylenediamine as a quencher. The sample was imaged using the 63X oil immersion objective of Leica confocal microscope. All incubations were carried out at room temperature, unless mentioned otherwise.

### Yeast two-hybrid analysis

Constructs containing genes cloned in pGBT9 (*TRP1*) or pGAD424 (*LEU2* selection marker), as described above, were transformed into the PJ69-4A yeast strain (Takara). To monitor growth on selective dropout plates, equal numbers of cells were spotted, and incubated for 3 days at 28°C. Quantitative estimations of various binary interactions were evaluated through analyses of β-galactosidase activity [34]. The results presented here are the average of three independent experiments, with each experiment comprising three technical replicates per sample. A two-tailed, unpaired t-test was conducted using GraphPad Prism 5 to determine the statistical significance.

### RT-PCR

Total RNA from trophozoites and encysting trophozoites was extracted using TRIZOL (Invitrogen) according to the manufacturer’s protocol. cDNA was prepared from the total RNA using Reverse Transcriptase (Invitrogen). The PCR reaction was performed with primers (S1 Table) corresponding to the internal sequence of *glist1* using the following conditions: initial denaturation at 95°C for 5 min, 30 cycles of amplification (95°C for 1 min, 55°C for 1 min, 72°C for 1 min), followed by post extension at 72°C for 10 min.

### Bioinformatic analysis of GlIst1

The sequences of GlIst1, ScIst1, ScVps4, GlVps4b, and HsVps4B were obtained from UniProt. The predicted tertiary structures were obtained from Alpha-Fold. InterPro and Pfam were used for the domain analysis. S3 Table contains the list of Gene ID and UniProt ID of the proteins utilized in domain analysis and structure prediction.

### Live cell imaging

BY4742 cells expressing GFP-GlIst1 and ScIst1-RFP were grown in a synthetic dropout medium until the O.D reached 0.5-0.6. Cells were harvested by centrifugation at 5000 rpm for 3 min. The supernatant was discarded, and the pellet was resuspended in YCM. 2 μl of the cells were mounted on a slide, and the sample was imaged with the 63X oil immersion objective of Leica confocal microscope.

### LC-ESI-MS/MS

250 ml of confluent trophozoite culture was incubated on ice for 30 min prior to collection via centrifugation at 1000g for 10 min. The harvested cells were then washed with 1X PBS at 1000g for 10 min and were subsequently resuspended in 200 μl of lysis buffer comprising 50 mM Tris pH 8.0, 120 mM NaCl, 5 mM EDTA, 1% Triton X-100, and a protease cocktail inhibitor (Sigma P8215). Cells were disrupted via sonication and the lysate was obtained by centrifugation at 13000 rpm for 10 min. The lysate was subsequently lyophilized. The lyophilized protein was then resuspended in 20 μl of 100 mM NH_4_HCO_3_ (Hi-Media). The protein concentration of the lysate was determined using Bradford assay, and the sample for mass-spectrometry was prepared using 2.5 μg/μlprotein. Each sample was combined with 25 μl trifluoroethanol (SRL) and 1.25 μl 200 mM DTT (Roche) and incubated at 60°C for 1h. Subsequently, 5 μl of 200 mM iodoacetamide (Sigma) was added, followed by incubation in the dark for 90 min. Then 1.25 μl of 200 mM DTT was added to the sample and incubated for 1 h. The final step involved the addition of 219 μl water and 100 mM NH_4_HCO_3_ to the sample. Overnight digestion of the sample was performed using 2.5 μl of 1 μg/μl trypsin (Promega V528A) at 37°C. The next day, the samples were centrifuged at 13000 rpm for 10 min, and the clear supernatant was collected. Lastly, 1 μl of 0.1% formic acid was added to the supernatant prior to loading the sample for LC-ESI-MS/MS (Waters Corporation). The data analysis was done by using three biological replicates. Progensis QI software was utilized for data acquisition and processing during LC-MS/MS analysis. The mass spectra of eluted peptides were recorded by the software. To identify the peptides, database searches were conducted against known protein sequence information from the UniProt database. In the mapping process, only peptide fragments that occurred at least three times were considered. The protein sequence coverage of GlIst1 was 55.8%.

## Results

### Sub-cellular distribution of the GlVps4 paralogs in trophozoites and during encystation

In previous reports, we have documented the distribution of GlVps4a and GlVps46a using polyclonal antibodies raised against these proteins [25, 31]. In trophozoites, GlVps4a predominantly localizes to the PVs located both along the cell periphery and around the bare zone surrounding basal bodies [25]. To further investigate its cellular localization under encysting conditions, we examined encysting trophozoites, both at 16 h and 48 post-induction (Fig 1A). At 16 h, GlVps4a was present in the PVs, but a minor pool of the protein was also detected in the cytoplasmic axonemes of the anterior flagella (AF) (Fig 1A). With the progress of encystation (48 h), the intensity of the signal at the flagella axonemes increased, indicating that this protein may play a role in the change in flagellar function and/or morphology leading up to the retraction of the flagellar axonemes (Fig 1A). The presence of minor pools of GlVps4a at the axonemes of the caudal and posterolateral flagella (CF and PF) pair further supports the possibility that GlVps4a regulates flagellar function during encystation. The protein was also present in the marginal plate, a unique structure just above the AF (Fig 1A). It was also present along the edges of the flanges, extending from both sides of the cell. The extension of the flange membranes is one of the key features of encysting trophozoites, which contributes to changes in cell morphology from the pear-shaped trophozoite to the ovoid cyst [35]. Since the ESCRT machinery is known to induce membrane curvature, the localization of GlVps4a along the deforming membrane during encystations may contribute to the membrane sculpting needed for the change in cellular morphology.

**Fig 1.**
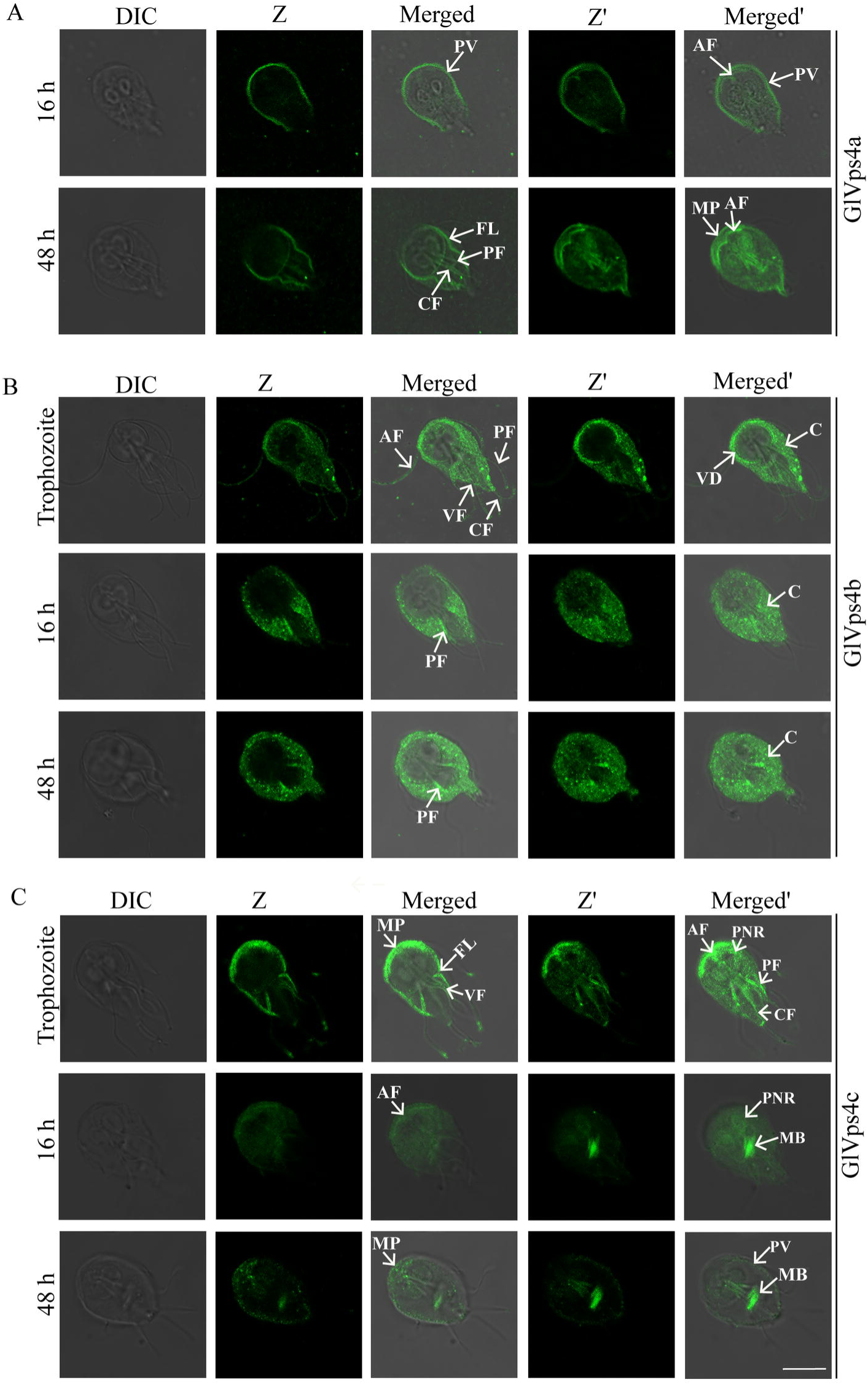
Sub-cellular distribution of the paralogs of GlVps4 in trophozoites and after induction of encystation. (A) Immunolocalization of GlVps4a at 16 h (upper panel) and 48 h (lower panel) post-induction of encystation. (B) Immunolocalization of GlVps4b in trophozoites (upper panel), 16 h (middle panel), and 48 h (lower panel) post-induction of encystations. (C) Immunolocalization of GlVps4c in trophozoites (upper panel), 16 h (middle panel), and 48 h (lower panel) post-induction of encystations. Arrows indicate the following cellular structures: PV: Peripheral Vesicles; AF: Anterior Flagella; PF: Posterior Flagella; VF: Ventral Flagella; CF: Caudal Flagella; VD: Ventral Disc; C: Cytosol; MB: Median Body; MP: Marginal Plate; FL: Flange; PNR: Perinuclear Region. Scale bar: 8 µm.

Next, we examined the cellular distributions of GlVps4b (Fig 1B) and GlVps4c (Fig 1C). In trophozoites, besides a cytoplasmic pool, GlVps4b was also present at the periphery of the VD andin the membrane-enclosed axonemes of all four types of flagella. It may be noted that unlike GlVsp4a, GlVps4b is absent at the PVs while its localization to the cytoplasm is consistent with the distribution of Vps4 ortholog in humans [36]. The presence of the protein at the VD periphery and flagella indicates that it may be required to maintain these cellular structures that are vital for *Giardia*’s virulence as these enable survival of the parasite within the host gut [37, 38]. Upon induction of encystation, while the cytoplasmic signal for Vps4b persisted, it was absent from the VD periphery or the membrane-encased flagella axonemes. Instead, it was observed at the cytoplasmic axonemes of the PF, where the signal intensified with the progress of encystation (Fig 1B, compare 16 h and 48 h). The observed redistribution of the signal from the flagella and the ventral disc during encystation raises the possibility that the loss of GlVps4b may precede the disassembly of these structures.

Curiously, the distribution of GlVps4c was the opposite of GlVps4b in association with the flagella axonemes. In trophozoites, GlVps4c was present in the cytoplasmic axonemes of the AF, PF, and CF (Fig 1C). Its signal was also detected at the membrane-bound axonemes of the ventral flagella (VF), PF and CF. Notably, signal enrichment was observed at the tips of all four types of flagella. The protein was also detected at the marginal plate and the perinuclear region. 16 h after induction of encystation, the signal at the cytoplasmic axonemes was lost, except for the AF, where the signal intensity was decreased compared to trophozoites. The major pool of the protein was at the median body, which is another microtubule-based structure in *Giardia* and is considered to be a reservoir of cytoskeletal elements [39]. The signal at the perinuclear region persisted at 16 h but was significantly diminished 48 h after encystation induction (Fig 1C). At this later time, while the signal at the median body persisted, the protein was present in punctate structures at the marginal plate region, and there was also a minor pool at the PVs. Considering the diverse sub-cellular distribution of the three paralogs of GlVps4, it may be concluded that they are likely to perform non-overlapping functions within the cell. In addition, they all localize to either microtubule-rich structures or at the sites of membrane deformation.

### Sub-cellular distribution of the GlVps46 paralogs in trophozoites and during encystation

Next, we investigated the distribution of two Vps46 paralogs, GlVps46a and GlVps46b. Quantitative PCR revealed that both paralogs of GlVps46 were expressed in trophozoites and at 16 and 48 h of encysting trophozoites (data not shown). The sub-cellular distribution of these paralogs, which share 75.8% identity and 85.8% similarity, indicated that while there was some overlap in their distribution, each was also localized to unique regions. Both paralogs were detected in the PVs, cytoplasm, VD periphery and the flagella axonemes. For GlVps46a, the cytoplasmic signal diminished considerably after 16 h of encystation when more cytoplasmic puncta were observed (Fig 2A). GlVps46a was also enriched in the ventrolateral flanges in trophozoites; the signal was also detected at this location at 16 h and 48 h post-induction of encystation. During encystation, the protein was also present as small puncta along the membrane-bound axonemes of the posterior flagella (PF) and CF. The signal at the VD periphery became more intense at 48 h encysting trophozoites and along the flanges (Fig 2A). For GlVps46b, in addition to the distribution mentioned above, it was enriched in the flagellar pores of the AF and PF and similar to GlVps4c, it was also observed at the flagellar tips (Fig 2B). Incidentally, none of the other ESCRT components we have studied displayed such a distribution at the flagella pores. During encystation, while the axoneme signal persisted at 16 h encysting cells, the pool of GlVps46b at the flagellar tip and the flagellar pores was lost (Fig 2B). At 48 h, the axoneme signal was lost. Based on these observations, we conclude that GlVps46b is likely to be involved in flagellar maintenance, and its flagellar pools recede later during encystation when flagella function is no longer needed. Our observations are consistent with a previous report showing that knockdown of CHMP4B (orthologous to Snf7 of yeast) causes defects in ciliary assembly and disintegration of cilia [40].

**Fig 2.**
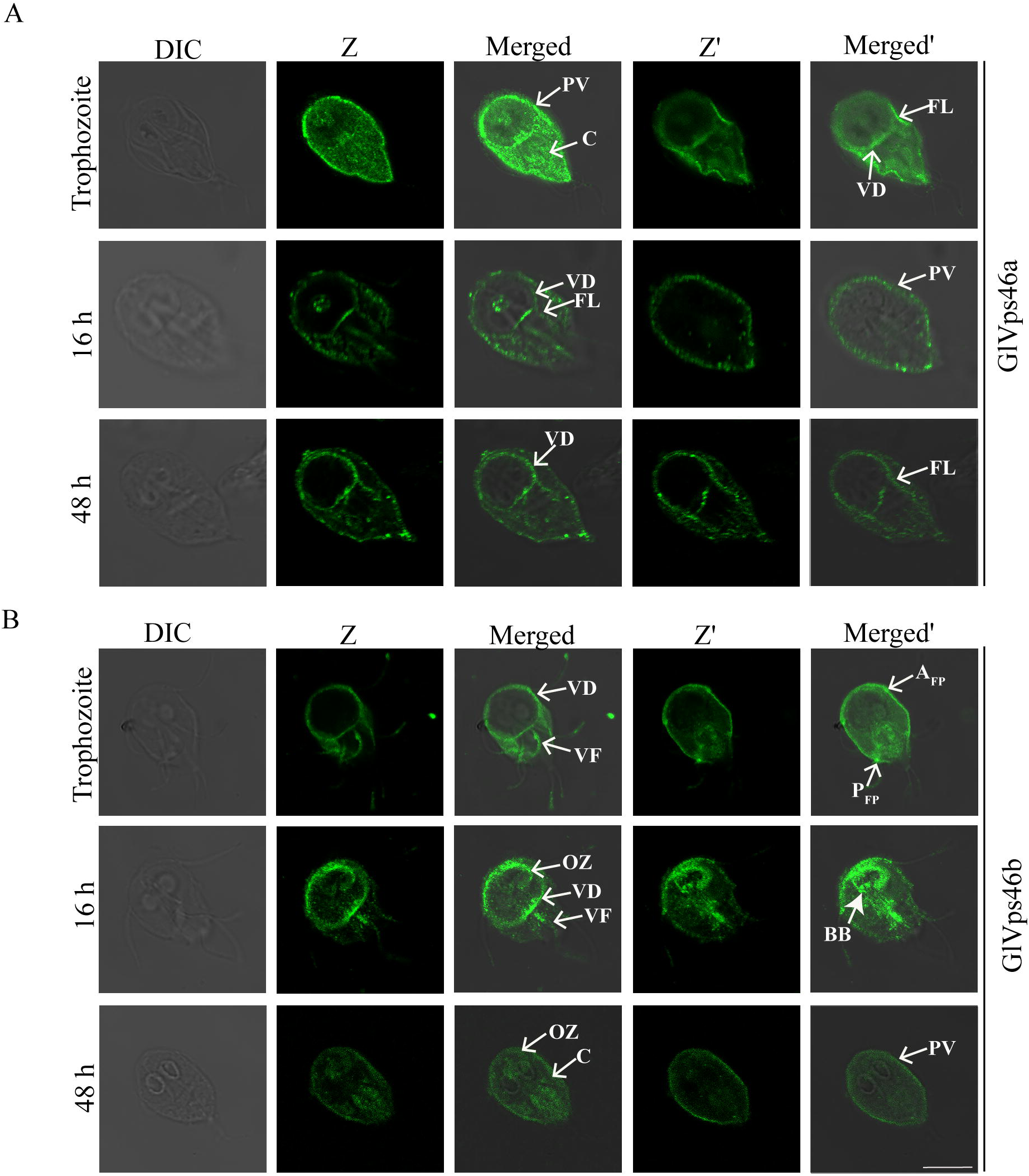
Sub-cellular distribution of the paralogs of GlVps46 in trophozoites and after induction of encystation. (A) Immunolocalization of GlVps46a in trophozoites (upper panel), 16 h (middle panel) and 48 h (lower panel) post-induction of encystations. (B) Immunolocalization of GlVps46b in trophozoites (upper panel), 16 h (middle panel) and 48 h post-induction of encystation (lower panel). Arrows indicate the following cellular structures: PV: Peripheral vesicles; C: Cytosol; AFP: Anterior Flagellar Pore; PFP: Posterior Flagellar Pore; FL: Flange; VD: Ventral Disc; VF: Ventral Flagella; OZ: Overlap Zone; BB: Basal Body. Scale bar: 8 µm.

Both GlVps46a and GlVps46b were present along the edges of the VD, but their distribution pattern during encystation was not identical; while the signal for GlVps46a increased with the progress of encystation (Fig 2A), that of GlVps46b was detected at the VD periphery of trophozoites and 16 h trophozoites but was absent at 48 h encysting trophozoites (Fig 2B). The VD is important for the survival of this parasite within the host gut [41]. The structure of this appendage involves a tight wrapping of the plasma membrane along its margins. Such sharp membrane bending involves the induction of a negative membrane curvature (bending of the membrane away from cytoplasm), and presently, only the ESCRT machinery is known to be capable of such membrane bending. GlVps46b was also detected in the overlap zone of the VD in trophozoites and the signal at this location was more pronounced in 16 h encysting cells (Fig 2B). The signal intensity diminished considerably, but was still evident in the 48 h encysting cells. Curiously, a substantial pool of GlVps46b was also present in the basal body at 16 h of encysting trophozoites. Based on the observations described above, we conclude that both the paralogs of this protein appear to play important roles in sustaining the unique cellular features that define this organism. In particular, they localize to sites where membranes bend sharply, such as the tips of flagella, flanges and VD periphery. Their re-localization during encystation also underscores their importance in bringing about morphological transitions during encystation.

The distribution of GlVps4 and GlVps46 paralogs in trophozoites and encysting trophozoites are summarized in S4 Table. The results of the localization study showed that on various occasions, the paralogous proteins Vps4 and Vps46 of *G. lamblia* were detected in the same sub-cellular location. However, there are also locations where these two sets of paralogs are present alone.

### Presence of an Ist1 ortholog in *Giardia lamblia*

Studies in yeast have shown that Vps46 regulates the recruitment of Vps4 to the endosomal surface [42]. We wanted to determine if there is a selective interaction between the paralogs of GlVps4 and GlVps46. Direct contact between ScVps46 and ScVps4 is well documented in yeast, and this interaction can be detected using the yeast two-hybrid (Y2H) assay [43]. Using the same assay, we failed to detect any interaction between the two paralogs of GlVps46 and the three paralogs of GlVps4, even though the yeast proteins showed high-affinity interaction (S3 Fig). Since both Vps46 and Vps4 play important roles in ESCRT-mediated membrane deformation, we hypothesized that although the *Giardia* orthologs do not interact directly, they may be brought into close apposition onto the membrane surface via their interactions with other ESCRT components. Studies in yeast and metazoans have shown that Ist1, a key ESCRT-III auxiliary component, interacts with both Vps4 and Vps46 [44, 45]. A search of the *Giardia* reference genome (Assemblage A isolate WB) indicated the presence of a putative GlIst1 (Gl50803_11129). Such putative GlIst1 orthologs were also encoded in the genomes of Assemblage B isolate GS (GL50581_3698) and Assemblage E isolate P15 (GLP15_1481) (Fig 3A). The ORF is also present in the recently-sequenced genome of strain BAH15c1 of Assemblage B (QR46_3855). However, it was not detected in the other sequenced *Giardia* genomes. Using reverse transcription PCR, we confirmed the expression of *glist1* in both the trophozoite and cyst stages of Assemblage A isolate WB, indicating that GlIst1 is produced in *Giardia* (Fig 3B).

**Fig 3.**
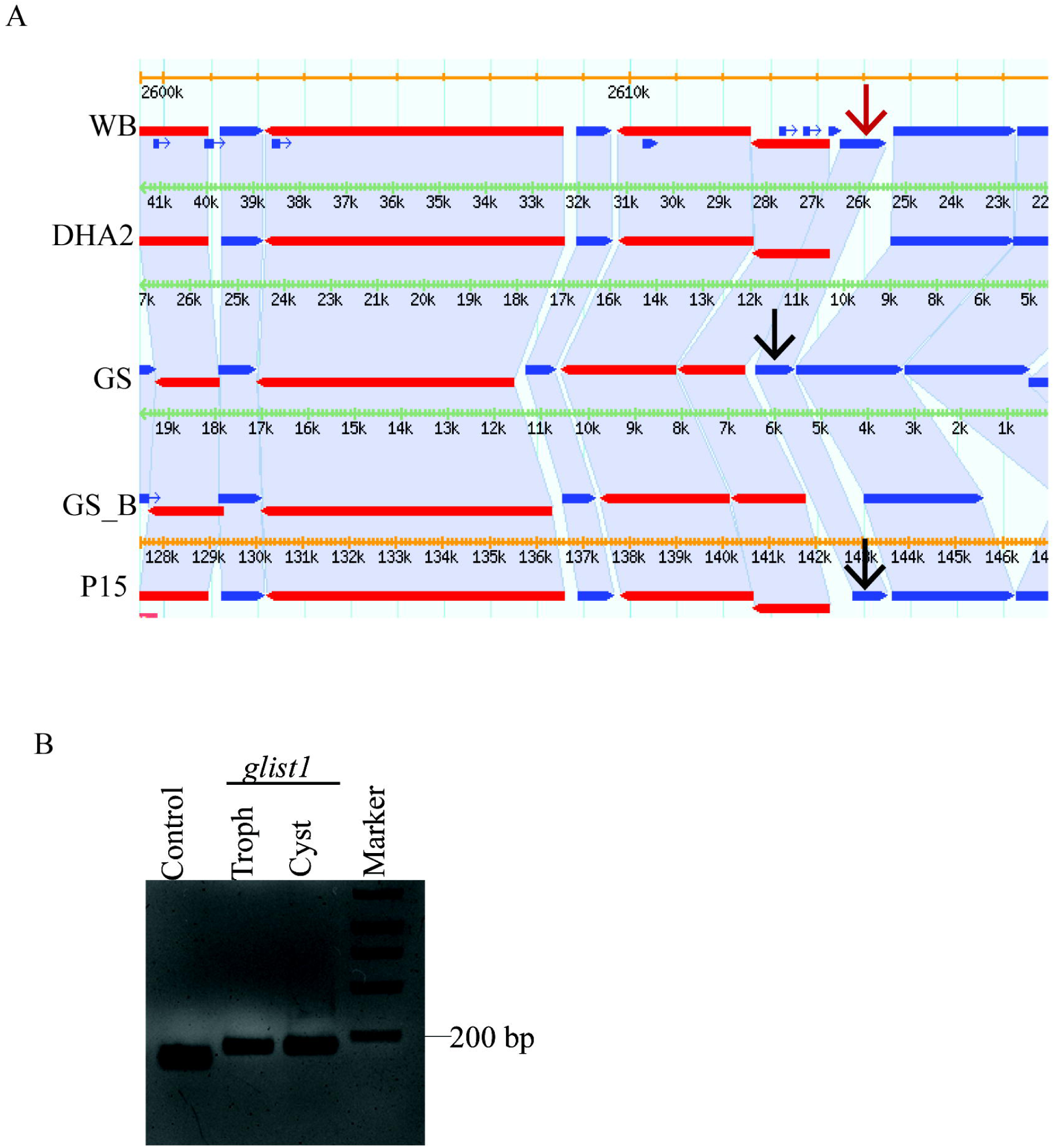
Synteny and expression of *glist1*. (A) Synteny of *glist1* in different isolates of *Giardia*. The red arrow indicates the location of *glist1* in the WB isolate, while the black arrows mark its position in other *Giardia* isolates. (B) Reverse transcription PCR to detect the expression of *glist1* in trophozoites and cysts. Expression of *glvps46b* serves as a positive control

### GlIst1-GlVps4b interaction is mediated by the MIM and MIT domain

If GlIst1 is functionally analogous to the yeast ortholog, it is likely to interact with some or all of the paralogs of GlVps4 and/or GlVps46. Using the Y2H assay, we first determined whether GlIst1 could physically interact with GlVps4 paralogs. The known interaction between the yeast orthologs, ScIst1 and ScVps4, served as the positive control. Based on the quantitative estimation of β-galactosidase activity, we also, observed that the expression of the LacZ reporter gene was turned on when cells co-expressed the Gal4 DNA binding domain fused to ScIst1 (BD-ScIst1) and the Gal4 activation domain fused with ScVps4 (AD-ScVps4) (Fig 4A). We detected β-galactosidase activity when BD-GlIst1 was co-expressed with AD-GlVps4b, but not when the former was co-expressed with either AD-GlVps4a or AD-GlVps4c, indicating selective binary interaction between GlIst1 and GlVps4b. A comparison of β-galactosidase activity indicated that this interaction within this *Giardia* protein pair was stronger than that between the yeast orthologs BD-ScIst1 and AD-ScVps4 (Fig 4A). Additionally, there was no interaction between BD-ScIst1 and AD-GlVps4b or between BD-GlIst1 and AD-ScVps4, indicating that Ist1 and Vps4 orthologs only interact with proteins from the same organism. Since the *Giardia* orthologs of both these proteins significantly diverge from their yeast counterparts, it is possible that the nature of the interaction between GlIst1 and GlVps4b may be different from that in yeast.

**Fig 4.**
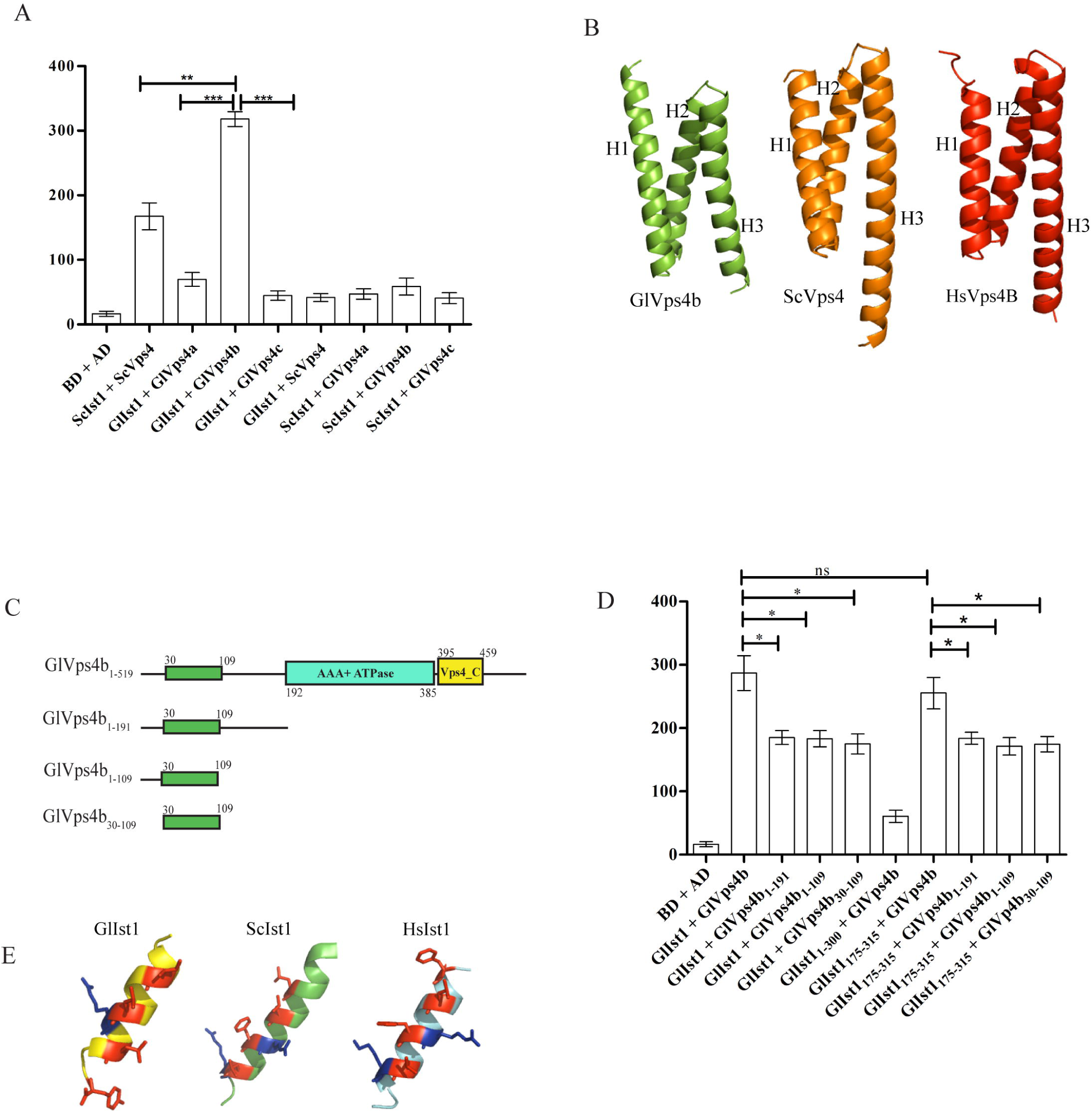
MIT domain of GlVps4b interacts with GlIst1. (A) β-galactosidase activity of PJ69-4A cells transformed with different constructs expressing fusion proteins featuring either the Gal4 DNA binding domain (BD) or its activation domain (AD). The expression of BD and AD alone served as negative control. (B) Alpha-Fold predicted secondary structure of GlVps4b, ScVps4 and HsVps4B. H1: Helix 1, H2: Helix 2 and H3: Helix 3 of the predicted three-helix bundle comprising the MIT domain. (C) Domain architecture of full-length GlVps4b or its different fragments used in the assay. (D) β-galactosidase activity of PJ69-4A cells transformed with different constructs expressing full-length or truncated versions of GlVps4b and GlIst1. Statistical significance: ns-not significant (p > 0.05), *-significant (p ≤ 0.05), **-very significant (p ≤ 0.01), and ***-extremely significant (p ≤ 0.001). (E) Alpha-Fold predicted secondary structure of the putative MIM of GlIst1 and the canonical MIM structural elements of ScIst1 and HsIst1.

Analyses of the interaction between Vps4 and Ist1 orthologs from multiple organisms show that it is mediated by the N-terminally-located MIT domain of Vps4 and the C-terminal MIT Interacting Motif (MIM) of Ist1 [46]. This interaction occurs via MIM1, wherein the MIM alpha helix binds to the groove formed by the second and third alpha helices of MIT domain [47]. We analyzed the sequence of GlVps4b and GlIst1 to determine if they have MIT and MIM, respectively. Interestingly, while the sequence of GlVps4b is 50.2% similar to ScVps4, none of the domain prediction databases hosted by InterPro recognized an MIT domain towards the N-terminus of the *Giardia* protein, even though the presence of the AAA ATPase domain and the Vps4_C domain was recognized. However, an MIT-like domain must be present in GlVps4b as it can functionally substitute ScVps4 [25]. Consistently, Alpha-Fold indicates the presence of a three-helix structure at its N-terminus that is structurally similar to the MIT domains of ScVps4 and VPS4B in humans (Fig 4B). These three helices span residues 30 to 110 of the protein. Based on this, we hypothesized that a non-canonical MIT domain is present at the N-terminus of GlVps4b. To test whether this three-helix bundle functions like the MIT domain of GlVps4b, we carried out progressive truncations of GlVps4b (Fig 4C). First, we compared the interactions of GlIst1 with either the full-length GlVps4b or the N-terminal portion of the protein, GlVps4b_1-191_, which includes all residues prior to the AAA ATPase domain. This region contains a short alpha helix near the N-terminus, followed by a three-helix bundle, a largely unstructured region and finally another short alpha helix. We used β-galactosidase activity to quantitatively compare these two interactions, and the results indicated that while there was some reduction in the affinity between the GlVps4b_1-191_ fragment and GlIst1, the decrease was not substantial as the p-value was <0.05 (Fig 4D). We carried out further truncations from the C-terminal end so that GlVps4b_1-109_ retains only up to the three-helix bundle. The β-galactosidase activity was nearly identical when either GlVps4b_1-191_ or GlVps4b_1-109_ interacted with GlIst1 (Fig 4D). Further truncation from the N-terminus generated the fragment GlVps4b_30-109_ that retained only the triple helix bundle. Even this fragment exhibited an almost similar affinity for GlIst1 as indicated by β-galactosidase activity which was similar to those of the previous two fragments (Fig 4D). These results indicate that residues spanning 30 to 109 of GlVps4b are sufficient to mediate the interaction between this GlVps4 paralog and GlIst1. Also, as these residues are predicted to form a three-helix bundle, GlVps4b likely contains a non-canonical MIT domain that cannot be recognized by any of the commonly used domain prediction databases.

Since GlVps4b’s interaction with GlIst1 is mediated via an MIT domain, this domain likely interacts with a MIM in GlIst1. The Alpha-Fold predicted the presence of a putative MIM-like amphipathic helix from residues 302 to 315 (Fig 4E). Similar to the canonical MIMs in the yeast and human orthologs, this helix is located near the C-terminal end of GlIst1, and it contains both hydrophobic (I305, L309, L312, and Y315) and positively charged (R310) residues (Fig 4E). Although Y315 is located in an unstructured region very close to the putative alpha helix, it may be induced into adopting an alpha-helical conformation upon binding to GlVps4b. To assess whether this region mediates the interaction with GlVps4b, we generated a truncation mutant, GlIst1_1-300_, fused with BD. Unlike full-length GlIst1, this mutant did not interact with AD-GlVps4b as evidenced by its β-galactosidase activity (Fig 4D). Notably, the interaction was restored when the C-terminal portion of GlIst1, GlIst1_175-315_, was expressed as indicated by β-galactosidase levels that were almost equal to those cells expressing the full-length GlIst1 and AD-GlVsp4b (Fig 4D). The interactions of GlIst1_175-315_ with those fragments of GlVps4b mentioned above were almost similar to that of the full-length GlIst1. Taken together, it may be inferred that despite the absence of the canonical MIT and MIM in GlVps4b and GlIst1, respectively, the MIT-MIM interaction appears to have been retained between GlIst1 and GlVps4b in this parasite.

### GlIst1 interacts only with GlVps46b

Next, we probed if GlIst1 can interact with the GlVps46 paralogs. β-galactosidase activity indicated that there is a strong interaction between BD-ScIst1 and AD-ScVps46, which is consistent with previous reports [48] (Fig 5A). Of the two Vps46 paralogs of *Giardia*, only GlVps46b could interact with GlIst1, and the quantification of the β-galactosidase activity indicates that the affinity between this *Giardia* protein pair was comparable to that observed for the orthologous protein pair from yeast. Similar to the species specificity observed in the case of the Ist1-Vps4 interaction described previously, here, too, we observed that there was no interaction between ScIst1 and GlVps46b or GlIst1 and ScVps46 (Fig 5A). This again underscores the divergence of the late-ESCRT proteins of *Giardia* from those of yeast.

**Fig 5.**
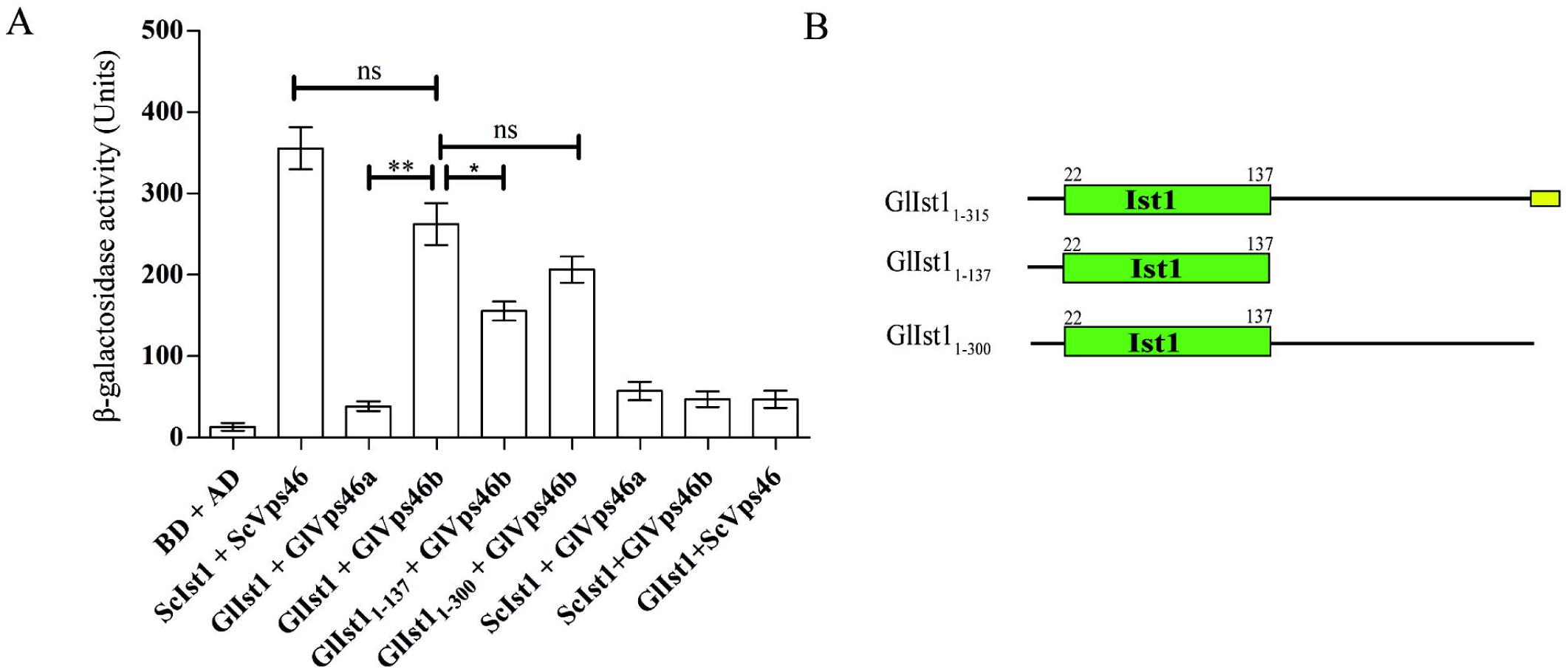
Ist1 domain of GlIst1 interacts with GlVps46b. (A) β-galactosidase activity of PJ69-4A cells transformed with different constructs expressing full-length or truncated versions of GlVps46b and GlIst1 fusion proteins featuring either the Gal4 DNA binding domain (BD) or its activation domain (AD). The expression of BD and AD alone served as negative control. (B) Domain architecture of full-length GlIst1 or its different fragments used in the assay. The Ist1 domain is depicted in green, and the MIM, spanning residues 302 to 315, is shown in yellow.

The interaction between Vps46 and Ist1 in higher organisms, including yeast and humans, is mediated by the N-terminal region of Ist1, which includes the Ist1 domain, and the MIM located at the C-terminus of Vps46 [44]. Sequence analysis revealed that GlIst1 shares a low level of similarity with its yeast and human orthologs, at 19.2% and 27.7%, respectively. Despite these low values, domain analyses revealed the presence of an Ist1 domain, spanning residues 22 to 137, in GlIst1 (Fig 5B). This 116 residues-long predicted Ist1 domain is considerably smaller than the same domain in the yeast and human orthologs, which are 164 and 171 residues, respectively. Hence, we were curious to determine if this smaller Ist1 domain can mediate the interaction with GlVps46b. While the β-galactosidase activity of transformants co-expressing BD-GlIst1_1-137_ and AD-GlVps46b was significantly higher than the corresponding negative control, it was lower than that observed with the full-length GlIst1 (Fig 5A). This suggests that additional segments of the GlIst1 protein are likely to contribute to the interaction with GlVps46b. Consistent with this hypothesis, a previous report suggested, that the α-5 of ScIst1 provides a binding surface for the MIM of ScVps46 [44]. Alpha-Fold indicates the presence of such a helix outside of the predicted Ist1 domain in GlIst1, spanning residues 172 to 187. To confirm that sequences beyond the predicted Ist1 domain are required for GlVps46b binding, we created a truncated version of this protein that is missing only the putative MIM helix, GlIst1_1-300_ (Fig 5B). Consistent with our expectation, this larger fragment interacted more efficiently with GlVps46b as the level of β-galactosidase was comparable to that produced with the full-length BD-GlIst1. Taken together, the above results indicate that GlIst1 can selectively interact with GlVps46b alone and this interaction is similar to that observed between the corresponding yeast orthologs. However, the interaction surfaces are likely to have undergone evolutionary changes that make each interaction species-specific.

### Membrane recruitment of GlIst1

In yeast, Ist1 is recruited to the endosomal membrane in a Vps46-dependent manner, after which it regulates Vps4 recruitment and activation [49]. Since the interaction dynamics between these three proteins are different in *Giardia* compared to yeast, we wanted to investigate if there are any differences in the functioning of GlIst1 *vis-à-vis* ScIst1. Towards this, we analyzed the post-translational modification profile of GlIst1. ScIst1 is known to undergo phosphorylation at multiple sites and some of these are documented to regulate its function [50]. When trophozoite extracts were subjected to LC-ESI-MS/MS analysis, we observed a unique myristoylation at an internal K43 residue. This modification was also evident in 16 h encysting trophozoites as well. There are no reports of either the yeast or human ortholog of Ist1 having such modification. The predicted Alpha-Fold structure of GlIst1 indicates that K43 is located in the loop region separating the first two helices (Fig 6A). It has been previously observed that upon membrane recruitment of Ist1, this region of the protein is in close proximity to the membrane surface [51], making it likely that a myristoyl group at this location will be able to interact with the membrane surface. Further, this post-translational modification is not in close proximity to the binding surfaces for either GlVps4b or GlVps46b. Hence it may not interfere with the binding of these two proteins with GlIst1.

**Fig 6.**
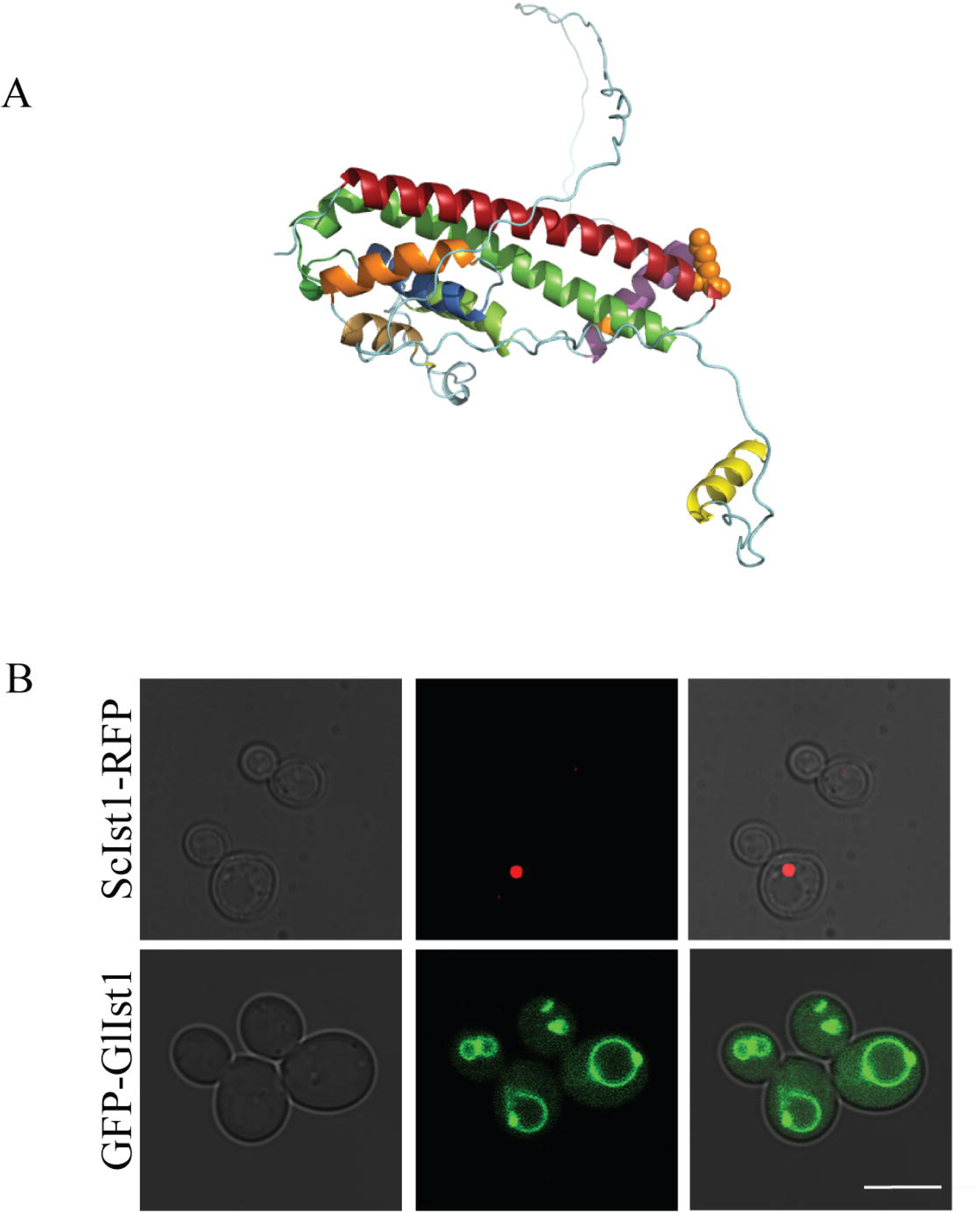
Difference in subcellular distribution of Ist1 orthologs in yeast and *Giardia*. (A) Position of the myristoylated K43 residue (orange spheres) in the Alpha-Fold structure of GlIst1. (B) Localization of GFP-GlIst1 and ScIst1-RFP proteins in yeast (BY4742). Scale bar: 8 µm.

The presence of a myristoyl group may affect the cellular distribution of GlIst1 compared to that of ScIst1, where no such modification is present. To test this, we localized fluorescently tagged ScIst1 and GlIst1 in yeast. The results indicate that while ScIst1 was exclusively endosomal, GlIst1 localized to both the endosome and the vacuole membrane. This difference in the distribution of the two orthologs indicates that GlIst1 has attributes that enable it to localize to the vacuole membrane, in addition to its localization at the endosome. This observation shows the functional divergence between the yeast and *Giardia* proteins and indicates that the late-ESCRT components of *Giardia* may have diverged from that of opisthokont model organisms such as yeast.

## Discussion

In previous studies, we documented the presence of three paralogs of GlVps4, two paralogs of GlVps46 and the presence of GlVps4a at the PV and the BZ of trophozoites and GlVps46a at the PVs. The present study is the first to elucidate the distribution of all the paralogs of GlVp4 and GlVps46 in both trophozoites and encysting trophozoites. This comprehensive immunolocalization data, obtained without the addition of any external tag that may interfere with incorporation into the ESCRT complex, provides new insight into the potential functions of the ESCRT components within the parasite. Based on the diverse sub-cellular locations to which these proteins are targeted, we believe that the ESCRT components play roles in multiple cellular processes in *Giardia*. Although some of these functions are similar to those documented in other species, some may have evolved specifically to address the challenges of sustaining the unique aspects of the biology of this gut parasite.

The paralogs of GlVps4 and GlVps46 are primarily present at the site of acute membrane bending, such as the periphery of the VD, edges of the flanges and tips of the flagella. All these structures are important virulence factors as they contribute greatly to the parasitic lifestyle of *Giardia* [52, 53]. VD is a key structural element that allows the parasite to remain attached to the host, thereby facilitating its survival. This study revealed the presence of certain ESCRT proteins, including Vps4b, and both Vps46a and Vps46b at the periphery of the VD, suggesting they may contribute to the parasite’s attachment. The flanges may mediate contact with the substratum and also drive shape change when a trophozoite changes to a cyst. Vps4a, Vps4c and Vps46a are found at the flange membrane, suggesting these ESCRT components may contribute towards trophozoite to cyst transition. Vps4c and Vps46b are also present at flagella tips. Studies in model organisms have indicated that ESCRTs are involved in the formation of such narrow membranous structures, such as the mid-body connecting the two daughter cells at the time of cytokinesis and the tubular extensions of endosomes that drive the recycling of various receptors. In the former case, ESCRT-III components are assembled inside the tubes while in the latter case, the ESCRT filaments form on the outside. Thus, ESCRT components at membrane protrusions along the edge of the flanges and at sites where there is wrapping of membranes around microtubule structures may be instrumental in bringing about the membrane deformation required at these sites. Incidentally, there appears to be some degree of functional segregation in the Vps46 paralogs. Vps46a is largely confined to sites of acute membrane bending (Fig 2A), while Vps46b also associates with microtubule structures, such as the OZ, the flagella axonemes and the flagella pores (Fig 2B). The Vps4 paralogs are also associated extensively with microtubule structures. All three of them are present at the cytoplasmic axonemes, even though at some locations, there is no overlap between them. For example, Vps4b is not present at the cytoplasmic axonemes of the anterior flagella and only Vps4c is present at the median body. However, we did not detect any such extensive association of the Vps46 paralogs with these microtubule structures. Such association of Vps4 paralogs may be due to the presence of the MIT domain in them. Whether Vps4 paralogs perform ESCRT-independent functions at these locations remains an open question as we cannot rule out the presence of minor pools of Vps46 paralogs at these locations. Such pools may be sufficient to discharge the specific function(s) for which Vps4 have been recruited.

*Giardia* has a unique cytoskeletal structure, primarily made of microtubules, and interestingly, various ESCRT components are observed in microtubule-rich regions such as flagella. It is also evident from the previous report that VPS4 is involved in the release of ectosomes from flagella in *Chlamydomonas* [54]. This study also found that paralogs of GlVps4 and GlVps46 are also observed at different parts of flagella, indicating their possible contribution to flagellar function.

The ESCRT components are very evident in PVs, which are involved in endocytosis, are crucial for transporting proteins and other molecules necessary for the parasite’s survival and pathogenicity, and help facilitate membrane remodeling during the internalization process [55]. Almost all ESCRT components are found in PVs, suggesting an ESCRT-dependent deformation of this compartment’s membrane. It has been previously reported that PVs have spherical and tubular shapes, which are associated with distinct functions [56]. Spherical PVs primarily serve as digestive compartments, whereas tubular or elongated PVs are primarily associated with dynamic exchange processes that facilitate the transport of materials between the PV lumen and the extracellular environment; their elongated shape may enhance their ability to interact with the plasma membrane. Similar endosomal tubule formation at the ER occurs with the help of CHMP1B, along with other ESCRTs and spastin. Our study also showed that Vps46b is present in PVs under various cellular conditions, suggesting that it may contribute to the formation of tubular PVs along with *Giardia*’s spastin.

Vps4, along with Vps46 and other ESCRT-III accessory components are responsible for the final stages of membrane sculpting and deformation. As previously reported, in higher eukaryotes Ist1 plays a key role in regulating the recruitment of Vps4, a critical step in ESCRT-mediated membrane sculpting. Our study revealed that GlIst1 interacts with both GlVps46 and GlVps4. It has been previously reported, ScIst1 interacts with ScVps4 and ScVps46. The question is, how does GlIst1 interact with GlVps46 and GlVps4? Does this resemble the Ist1-Vps4 interaction observed in other eukaryotes? In higher eukaryotes, the MIT domain of Vps4 in the N-terminal region interacts with the MIM motif of the ESCRT-III components present in the extreme C-terminal region. Our results support the presence of a similar MIT-MIM interaction between the *Giardia* orthologs, even though both the MIT of GlVps4b and the MIM of GlIst1 are non-canonical. Similarly, the interaction between GlIst1 and GlVps46b occurs via the N-terminal Ist1 domain of the former, and this mode of interaction also seems to be conserved, as in humans and yeast. This suggests that while sequence divergence exists between the late-ESCRT components of *G. lamblia* and the higher eukaryotic orthologs, the nature of the interaction appears to have been conserved during the evolutionary divergence of this parasite.

A previous study proposed that Ist1 could both inhibit and enhance Vps4 ATPase activity. This dual role is facilitated by different elements within Ist1, such as MIM and a conserved ELYC sequence. The ELYC region contributes to the negative regulation of Vps4 by Ist1 [49], whereas the MIM region is responsible for the activation of Vps4. In GlIst1 no such EYLC region was found, which may contribute to the differential interplay between Vps46, Vps4 and Ist1. In yeast, the binding of ScVps46 to ScIst1 causes a conformational change in the ScIst1 core. This structural alteration is believed to be essential for transforming ScIst1 from an inhibitor to a stimulator of ScVps4 ATPase activity [49]. Similarly, in *Giardia*, one pathway of the ESCRT machinery may function by GlIst1 being independently recruited to the membrane through myristoylation and binding to Vps46. This interaction may facilitate a conformational change in Ist1, promoting it’s binding to Vps4 and ultimately leading to the completion of the MVB pathway. Interestingly, although there are no reports of myristoylation of Ist1 in other eukaryotes, yeast Vps20 is recruited to the membrane by myristoylation at its N-terminal glycine residue [57]. Thus, throughout evolution, myristoylation as a means of membrane recruitment of ESCRT-III components may have arisen as an independent event.

## Acknowledgement

The authors acknowledge all members of the Sarkar Laboratory for discussions and valuable comments. DNA sequencing and confocal imaging were carried out at the CIF of Bose Institute. Prantik Saha and Sheolee Ghosh Chakraborty are acknowledged for assisting with confocal imaging. This study was supported by research funds to SS from the Bose Institute (R/16/19/1620).

## Supporting Information

**S1 Table.** Sequences of primers used in this study

**S2 Table.** List of constructs used in this study

**S3 Table.** List of Gene ID and UniProt ID used in this study

**S4 Table**. Summary of the localization of different paralogs of GlVps4 and GlVps46

**S1 Fig. Specificity of antibodies against the paralogs of GlVps4.**

(A) Cells treated with pre-immune sera collected from animals prior to immunization with GlVps4b (upper panel) or with GlVps4c (bottom panel). Scale bar 8 µm. (B) Western blotting of the three 6x-His tagged GlVps4 paralogs expressed in *E. coli* and then purified from the bacterial extracts. Blots were incubated with anti-GlVps4a antibody (left), with anti-GlVps4b antibody (middle), and with anti-GlVps4c antibody (right). The detections of bands of sizes ∼56kDa, ∼60 kDa, and ∼30 kDa demonstrate the specificity of GlVps4a, GlVps4b, and GlVps4c antibodies, respectively. (C) Western blots were performed using trophozoite extracts. The blot was developed using the anti-GlVps4b antibody (left), while the blot was developed using anti-GlVps4c (right).

**S2 Fig. Specificity of antibodies against the paralogs of GlVps46**. (A) Cells treated with pre-immune sera were collected from animals prior to immunization with GlVps46a (upper panel). Cells treated with pre-immune sera were collected from animals prior to immunization with GlVps46b (bottom panel). Scale bar 8 µm. (B) Western blot analysis was performed using PJ69-4A transformants expressing either BD-tagged GlVps46a or BD-tagged GlVps46b. The western blot was performed with anti-GlVps46b antibody (left) and anti-GlVps46a antibody (right). A∼37.4 kDa band (right) and a ∼37.8 kDa band (left) were detected. (C) Western blots were performed using trophozoite extracts. The blot was developed using the anti-GlVps46a antibody (left) and anti-GlVps46b antibody (right). All western blots were performed using 26616 (Thermo) markers except for GlVps46b (26619).

**S3 Fig. Lack of binary interaction between the paralogs of GlVps4 and GlVps46 using yeast two-hybrid assay.** PJ69-4A cells were transformed with various BD and AD fusion combinations, as indicated in the figure. Growth of the PJ694-A transformants were monitored on SD leu^-^trp^-^ (left panel), SD leu^-^trp^-^his^-^ with 2.5 mM 3-AT (middle panel), and leu^-^trp^-^ade^-^ (right panel). The experiment was repeated with constructs expressing proteins with reversal of BD or AD fusion (bottom panels).

**S4 Fig. Quantification of** β**-galactosidase activity in all PJ69-4A transformants.** (A) An extended representation of the data is presented in Fig 4A, (B) Fig 4D and (C) Fig 5A

**S5 Fig. Alpha-Fold predicted structures of Ist1 orthologs.** (A) GlIst1 of *Giardia lamblia*. (B) Ist1 of *S. cerevisiae*. The helices are colour-coded to indicate their position relative to the N-terminus: α-1 (red), α-2 (green), α-3 (blue), α-4 (orange), α-4a(sand), α-4b(chartreuse green) and α-5(magenta) and the MIM (yellow). The unstructured regions are in cyan.

**S6 Fig. Detection of myristoylationin GlIst1 using LC-MS/MS** (A) Peptide fragments of GlIst1 showing myristoylation. (B) Overall detection of peptide fragments is highlighted in yellow, with the myristoylated K residue highlighted in red. The peptide sequence coverage for GlIst1 was 55.8%

## Notes

### Competing Interest Statement

The authors have declared no competing interest.

