## Supplementary table 1 for "Divergent Functions of Late ESCRT Components in *Giardia lamblia*: Insights from Subcellular Distributions and Protein Interactions"

Sequences of primers used in this study

| **SI No.** | **Sequence 5’→3’** | **Purpose** |
| --- | --- | --- |
| 1 | TAGAATTCATGCACGGTCAGTTCTTTGCG | *glvps4b* N-ter _(1-109)_ Forward |
| 2 | ATTTGTCGACTTAGGCCTGAGCATCGTGTAT | *glvps4b* N-ter _(1-109)_ Reverse |
| 3 | GCGGGAATTCATGAATCAAGTTAGAAAAAAAATCGC | *glvps4b* N-ter _(30-109)_ Forward |
| 4 | TTGTCGACTTAGGCCTGAGCATCGTGTATCTGATCTAA | *glvps4b* N-ter _(30-109)_ Reverse |
| 5 | CAGAATTCATGCACGGTCAGTTCTTTGCG | *glvps4b* N-ter _(1-191)_ Forward |
| 6 | ATTTGTCGACTTACTCGCGCTTGCCTTGGAA | *glvps4b* N-ter _(1-191)_ Reverse |
| 7 | GCCGAATTCATGCCAAAGATTAATGAACTCG | *glist1* N-ter _(1-137)_ Forward |
| 8 | ATTGGATCCTCATAGTCTGAGGGCCTGGTC | *glist1* N-ter _(1-137)_ Reverse |
| 9 | GCGAATTCATGGATTTGGAGTACAACAAG | *glist1* C-ter _(175-315)_ Forward |
| 10 | TTGGATCCTCAGTAGTTGTTCAGCTT | *glist1* C-ter _(175-315)_ Reverse |
| 11 | GCGGAATTCATGCCAAAGATTAATGAAC | *glist1*_(1-300)_ Forward |
| 12 | ATTGGATCCTCATGTTAGGGGCTTGTCATC | *glist1*_(1-300)_ Reverse |
| 13 | GCGGAATTCATGCCAAAGATTAATGAAC | *glist1* Forward |
| 14 | ATAGGATCCGTAGTTGTTCAGCTTTCTG | *glist1* Reverse |
| 15 | GTTGGGAACCTGAAGGACC | *glist1* qPCR Forward |
| 16 | GATGGACCCGAACAGCTTGAC | *glist1* qPCR Reverse |
| 17 | GACCAGGCTATGGAGCTGAAG | *glvps46b* qPCR Forward |
| 18 | GAGGTTTGCCTTTGTGTTGGC | *glvps46b* qPCR Reverse |
| 19 | GCGAATTCATGGGGAAGATAGATCTTACC | *glvps46a* Forward |
| 20 | CGTGGATCCTTTGTCCGTTTACTCGACG | *glvps46a* Reverse |
| 21 | CCGAATTCATGGGCAAAGTGGATCTGG | *glvps46b* Forward |
| 22 | CCGGATCCTTACACCATCTCCGCCGCTC | *glvps46b* Reverse |
| 23 | CGGGATCCGTATGTCACGTAATTCTGCA | *ScVPS46* Forward |
| 24 | AATGTCGACTCAGCCCCTCAATGCTCTT | *ScVPS46* Reverse |
| 25 | CCGAATTCATGACCATTGTTCCTGGTCG | *glvps4a* Forward |
| 26 | TGGCGTCGACTATAACAGCTCAGGTGGAGC | *glvps4a* Reverse |
| 27 | CAGAATTCATGCACGGTCAGTTCTTTGC | *glvps4b* Forward |
| 28 | CTCGGTCGACGCATGGCTACATTAGGATC | *glvps4b* Reverse |
| 29 | CAGAATTCATGTGCTCGCGGCTCAACC | *glvps4c* Forward |
| 30 | TGGCGTCGACTGGCTCTATGAGTGCTTAAAC | *glvps4c* Reverse |
| 31 | CGGAATTCATGAGCACGGGAGATTTTTTAAC | *ScVPS4* Forward |
| 32 | CGGGATCCCTAGTTACCTTCTTGACCAAAATC | *ScVPS4* Reverse |
| 33 | CAGGATCCATCACAATGGCTCCGTCAATGATTC | *ScIST1* Forward |
| 34 | AGCCGTCGACTTATTTTCTGCGTAAAGCGTC | *ScIST1* Reverse |
| 35 | TTGGAATTCATGCACGGTCAGTTCTTTG | *glvps4b* Forward pHis-TEV |
| 36 | TTGCGTCGACTTACTTGGAAGTTTTTCC | *glvps4b* Reverse pHis-TEV |
| 37 | AAGGAATTCATGTGCTCGCGGCTCAA | *glvps4c* Forward pHis-TEV |
| 38 | CCCAAGCTTCTATCCACGATTGAGAGATA | *glvps4c* Reverse pHis-TEV |
| 39 | GGGGAATTCGGCAAAAAAATG | *glvps46a* Forward pET21d |
| 40 | CCCAAGCTTCCGTTTACTCG | *glvps46a* Reverse pET21d |
| 41 | ATAGGATCCATGGGCAAAGTGGATCTG | *glvps46b* Forward pHis-TEV |
| 42 | TTCAAGCTTTTACACCATCTCCGCCGC | *glvps46b* Reverse pHis-TEV |
| 43 | TGGGATCCATGGCTCCGTCAATGCTTC | *ScIST1* Forward for cloning  in pRS425-RFP |
| 44 | GCGTCGACTTTTCTGCGTAAAGCGTC | *ScIST1* Reverse for cloning in pRS425-RFP |
| 45 | CGTGAATTCCATGCCAAAGATTAATGAAC | *glist1* Forward for cloning  in pGOGFP426 |
| 46 | AGCCGTCGACTCAGTAGTTGTTCAGCTTTC | *glist1* Reverse for cloning  in pGOGFP426 |
