## Supplementary table 2 for "Divergent Functions of Late ESCRT Components in *Giardia lamblia*: Insights from Subcellular Distributions and Protein Interactions"

| Construct Name | Cloned gene | Description | Primers used |
| --- | --- | --- | --- |
| pNP1 | *ScVPS4* | p(ADH1)-GAL4AD-*ScVPS4*(pGAD424*LEU2*) | 31 and 32 |
| pNP2 | *ScVPS4* | p(ADH1)-GAL4BD-*ScVPS4*(pGBT9 *TRP1*) | 31 and 32 |
| pNP3 | *glvps4a* | p(ADH1)-GAL4AD-*glvps4a*(pGAD424 *LEU2*) | 25 and 26 |
| pNP4 | *glvps4a* | p(ADH1)-GAL4BD-*glvps4a*(pGBT9 *TRP1*) | 25 and 26 |
| pNP5 | *glvps4b* | p(ADH1)-GAL4AD-*glvps4b* (pGAD424 *LEU2*) | 27 and 28 |
| pNP6 | *glvps4b* | p(ADH1)-GAL4BD-*glvps4b* (pGBT9 *TRP1*) | 27 and 28 |
| pNP7 | *glvps4c* | p(ADH1)-GAL4AD-*glvps4c* (pGAD424 *LEU2*) | 29 and 30 |
| pNP8 | *glvps4c* | p(ADH1)-GAL4BD-*glvps4c* (pGBT9 *TRP1*) | 29 and 30 |
| pNP9 | *ScVPS46* | p(ADH1)-GAL4AD-*ScVPS46* (pGAD424 *LEU2*) | 23 and 24 |
| pNP10 | *ScVPS46* | p(ADH1)-GAL4BD-*ScVPS46*(pGBT9 *TRP1*) | 23 and 24 |
| pNP11 | *glvps46a* | p(ADH1)-GAL4AD-*glvps46a*(pGAD424 *LEU2*) | 19 and 20 |
| pNP12 | *glvps46a* | p(ADH1)-GAL4BD-*glvps46a* (pGBT9 *TRP1*) | 19 and 20 |
| pNP13 | *glvps46b* | p(ADH1)-GAL4AD-*glvps46b* (pGAD424 *LEU2*) | 21 and 22 |
| pNP14 | *glvps46b* | p(ADH1)-GAL4BD-*glvps46b* (pGBT9 *TRP1*) | 21 and 22 |
| pNP15 | *ScIST1* | p(ADH1)-GAL4BD-*ScIST1*(pGBT9 *TRP1*) | 33 and 34 |
| pNP16 | *glist1* | p(ADH1)-GAL4BD-*glist1*(pGBT9 *TRP1*) | 13 and 14 |
| pNP17 | *glist1*_1-300_ | p(ADH1)-GAL4BD-*glist1*_1-300_ (pGBT9 *TRP1*) | 11 and 12 |
| pNP18 | *glist1*_1-137_ | p(ADH1)-GAL4BD-*glist1*_1-137_ (pGBT9 *TRP1*) | 7 and 8 |
| pNP19 | *glist1*_175-315_ | p(ADH1)-GAL4BD-*glist1*_175-315_ (pGBT9 *TRP1*) | 9 and 10 |
| pNP20 | *glvps4b*_1-191_ | p(ADH1)-GAL4AD-*glvps4b*_1-191_ (pGAD424 *LEU2*) | 5 and 6 |
| pNP21 | *glvps4b*_1-109_ | p(ADH1)-GAL4AD-*glvps4b*_1-109_ (pGAD424 *LEU2*) | 1 and 2 |
| pNP22 | *glvps4b*_30-109_ | p(ADH1)-GAL4AD-*glvps4b*_30-109_ (pGAD424 *LEU2*) | 3 and 4 |
| pNP23 | *glvps4b* | p(T7)-HisTEV-*glvps4b* (pHis-TEV *Amp*) | 35 and 36 |
| pNP24 | *glvps4c* | p(T7)-HisTEV-*glvps4c* (pHis-TEV *Amp*) | 37 and 38 |
| pNP25 | *glvps46a* | p(T7)-His*glvps46a* (pET21d *Amp*) | 39 and 40 |
| pNP26 | *glvps46b* | p(T7)-HisTEV-*glvps46b* (pHis-TEV *Amp*) | 41 and 42 |
| pNP27 | *ScIST1*-RFP | p(PYK1)-S*cIST1*-RFP (pRS425-*LEU2*) | 43 and 44 |
| pNP28 | GFP-*glist1* | p(CPY)-GFP-*glist1* (pRS426-*URA3*) | 45 and 46 |

**Constructs used in this study**
