## Supplementary table 3 for "Divergent Functions of Late ESCRT Components in *Giardia lamblia*: Insights from Subcellular Distributions and Protein Interactions"

**List of Gene ID and UniProt ID used in this study**

| Gene name | Gene ID | Uniprot ID |
| --- | --- | --- |
| *glist1* | Gl50803_0011129 | A8BUY9 |
| *ScIST1* | YNL265C | P53843 |
| *ScVPS4* | YPR173C | P52917 |
| *glvps4b* | Gl50803_16795 | A8BUC0 |
| *HsVps4B* | 9525 | O75351 |
