## Supplementary table 4 for "Divergent Functions of Late ESCRT Components in *Giardia lamblia*: Insights from Subcellular Distributions and Protein Interactions"

| Components Components | Stage | PV | VD | Axoneme  M C | | FT | FP | FL | MP | PNR | BB | MB | C | BZ | OZ |
| --- | --- | --- | --- | --- | --- | --- | --- | --- | --- | --- | --- | --- | --- | --- | --- |
| GlVps4a  Gl50803_101906 | Troph | Yes | --- | --- | --- | --- | --- | --- | --- | --- | --- | --- | Yes | Yes | --- |
|  | 16 h | Yes | --- |  | Yes | --- | --- | --- | --- | --- | --- | --- | --- | --- | --- |
|  | 48 h | Yes | --- | --- | Yes  (AF) | --- | --- | Yes | Yes | --- | --- | --- | --- | --- | --- |
| GlVps4b  Gl50803_16795 | Troph | --- | Yes | Yes | Yes | --- | --- | --- | --- | --- | --- | --- | Yes | --- | --- |
|  | 16 h | --- | --- | --- | Yes | --- | --- | --- | --- | --- | --- | --- | Yes | --- | --- |
|  | 48 h |  | --- | --- | Yes | --- | --- | --- | --- |  | --- | --- | Yes | --- | --- |
| GlVps4c  Gl50803_15469 | Troph | Yes | --- | Yes | Yes | Yes | --- | Yes | Yes | Yes | --- | --- | --- | --- | --- |
|  | 16 h | --- | --- | --- | Yes | --- | --- | --- | --- | Yes | --- | Yes | Yes | --- | --- |
|  | 48 h | Yes | --- | --- | --- | --- | --- | --- | Puncted | Faint | --- | Yes | --- | --- | --- |
| GlVps46a  Gl50803_15472 | Troph | Yes | Yes | --- | --- | --- | --- | Yes | --- | --- | --- | --- | Yes | --- | --- |
|  | 16 h | Yes | Yes | Puncted | --- | --- | --- | Yes | --- | --- | --- | --- | Puncted | --- | --- |
|  | 48 h | Yes | Yes | --- | --- | --- | --- | Yes | --- | --- | --- | --- | Puncted | --- | --- |
| GlVps46b  Gl50803_24947 | Troph | Yes | Yes | Yes | --- | Yes | Yes | Faint | --- | --- | --- | --- | --- | --- | Yes |
|  | 16 h | Yes | Yes | Yes | --- | --- | --- | --- | --- | --- | Yes | --- | --- | --- | Yes |
|  | 48 h | Yes | --- | --- | --- | --- | --- | --- | --- | --- | --- | --- | Yes | --- | Yes |

**Cellular distribution of the ESCRT paralogs**

PV: Peripheral Vesicles; VD: Ventral Disc; PNR: Perinuclear Region; FL: Flange; Axoneme (C): Cytosolic Axoneme; Axoneme (M): Membrane-bound Axoneme; FT: Flagellar Tip; FP: Flagellar Pore; MP: Marginal Plate; BB: Basal Body; MB: Median Body; C: Cytosol; BZ: Bare Zone; OZ: Overlap Zone
