## Supplementary figures and images for "Divergent Functions of Late ESCRT Components in *Giardia lamblia*: Insights from Subcellular Distributions and Protein Interactions"

### Supplementary Figure 1

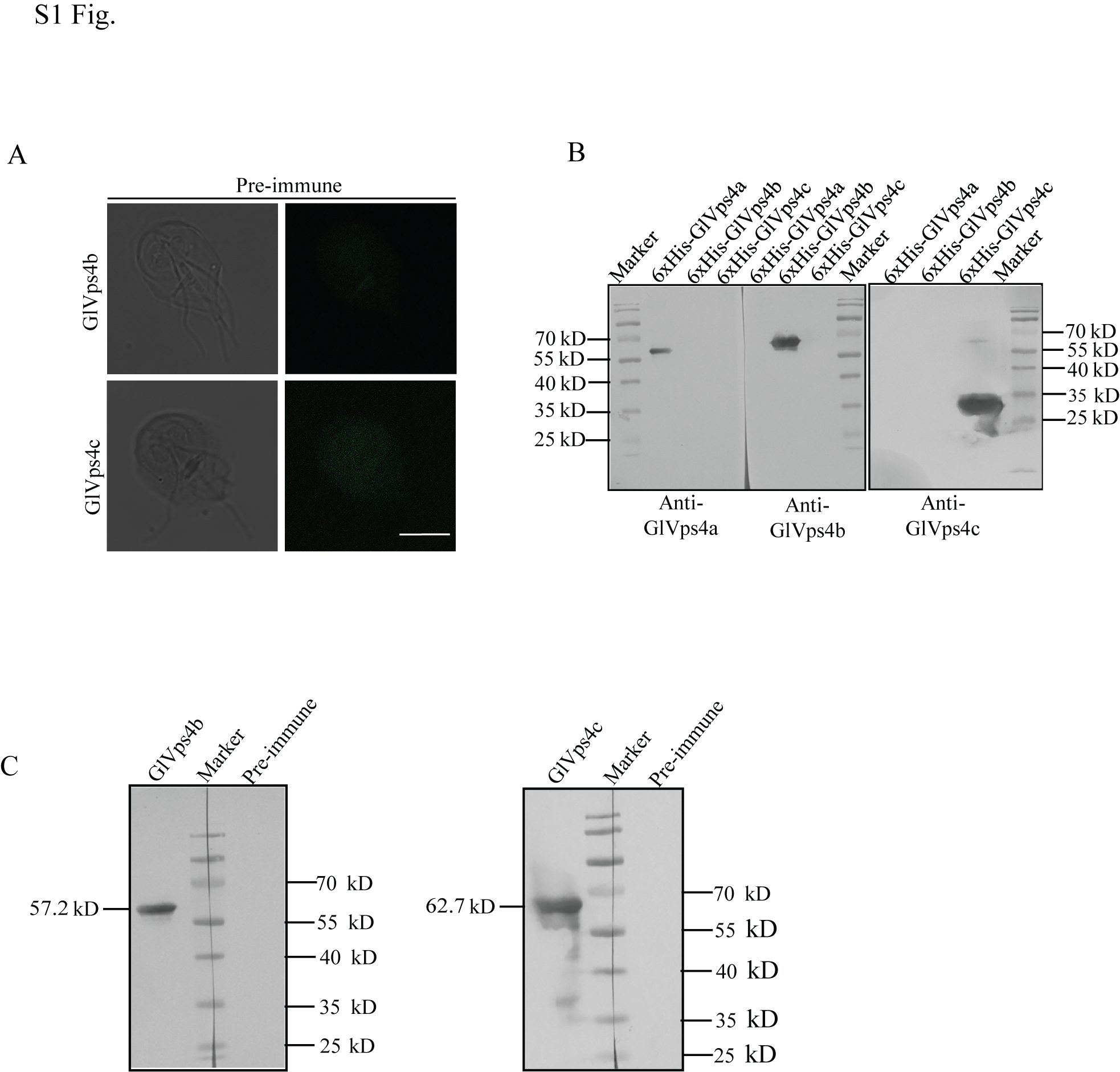

### Supplementary Figure 2

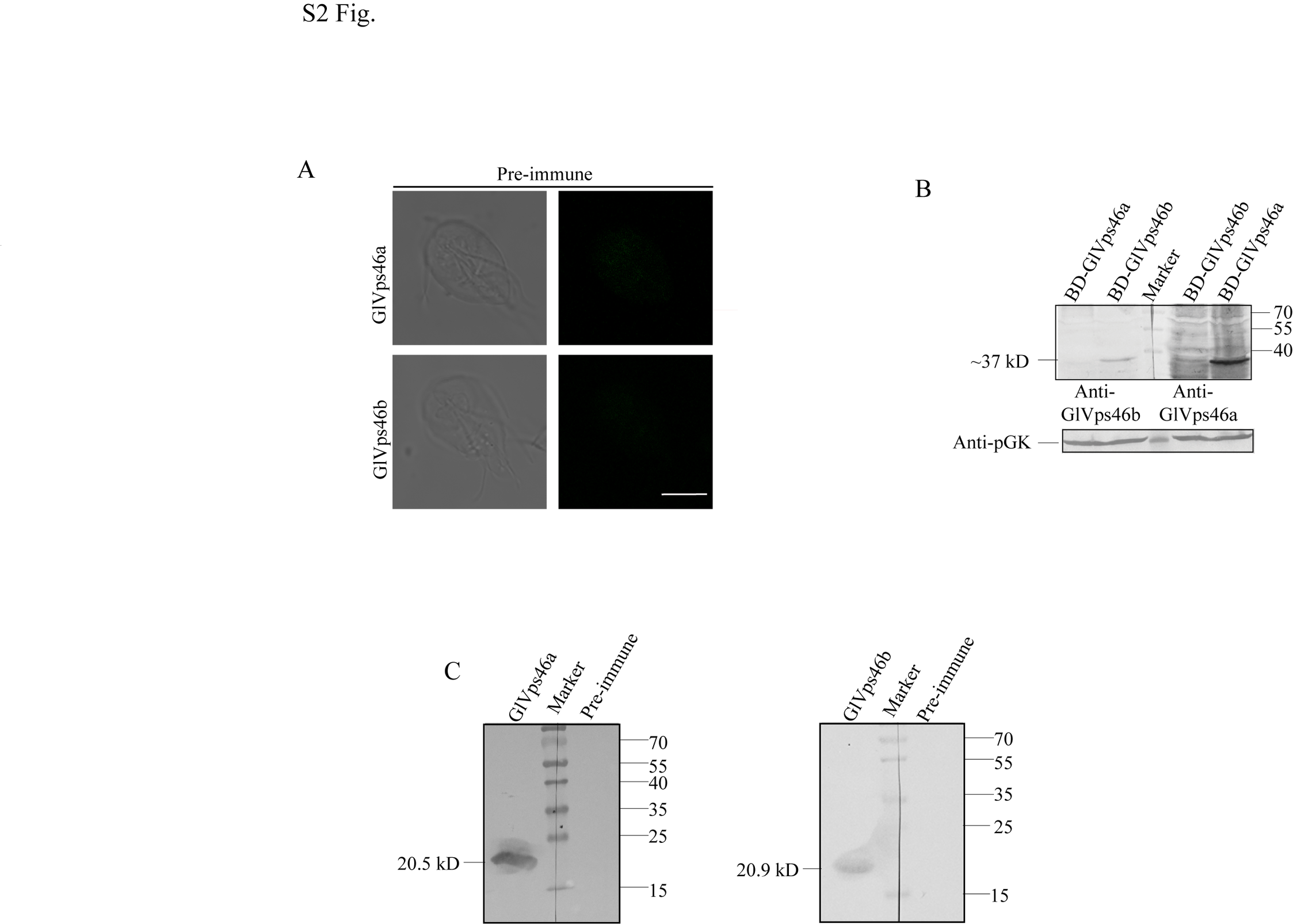

### Supplementary Figure 3

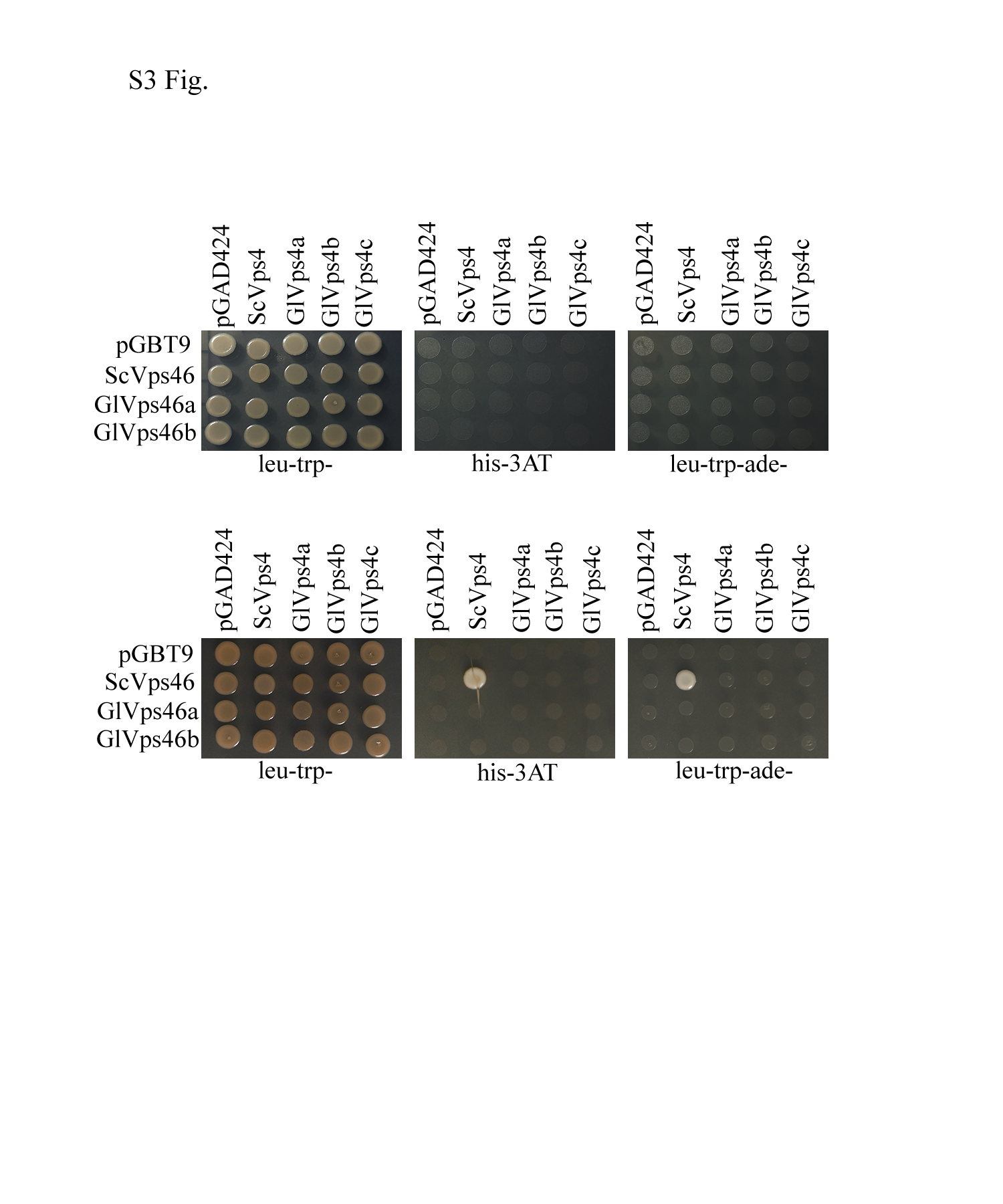

### Supplementary Figure 4

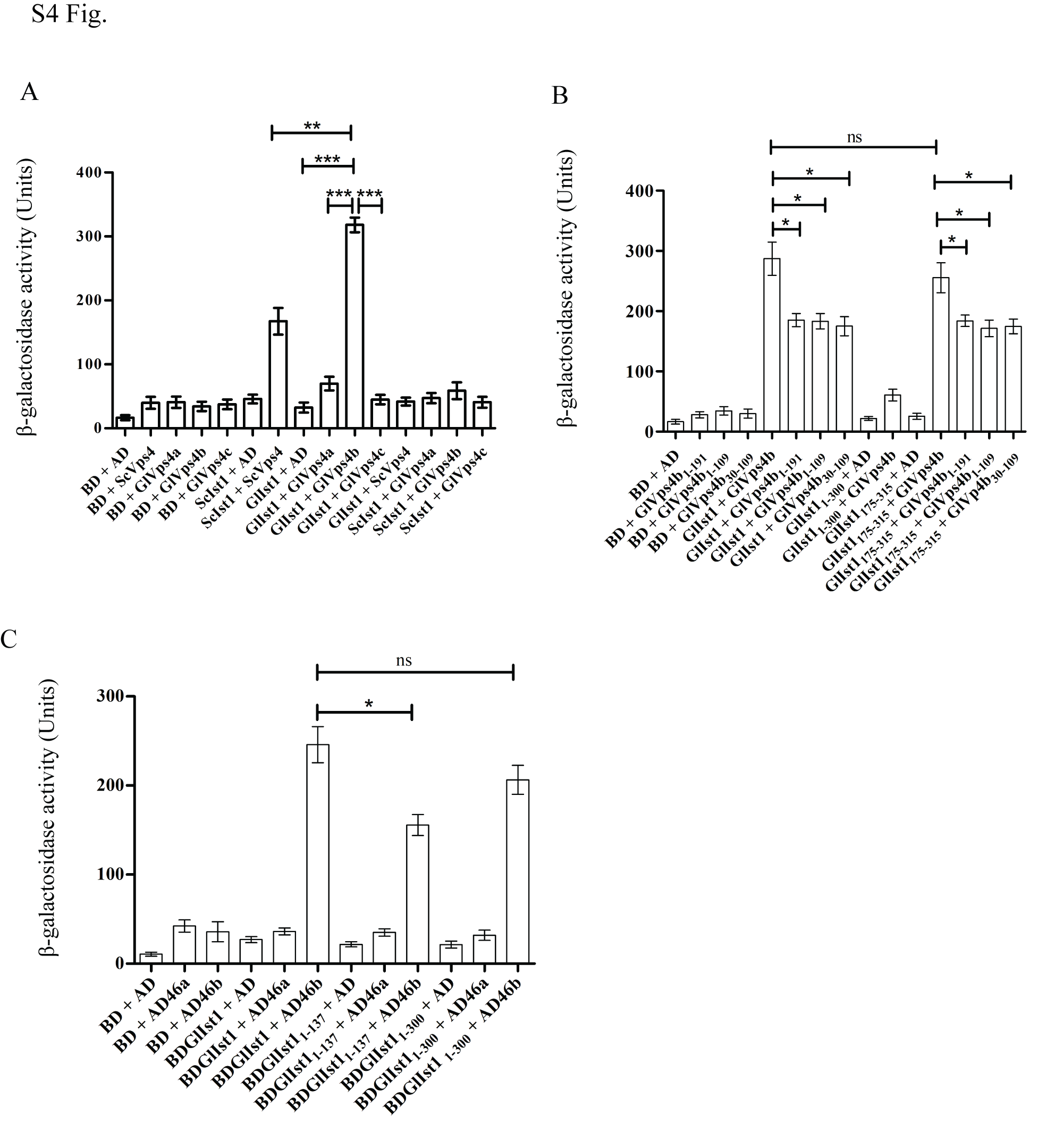

### Supplementary Figure 5

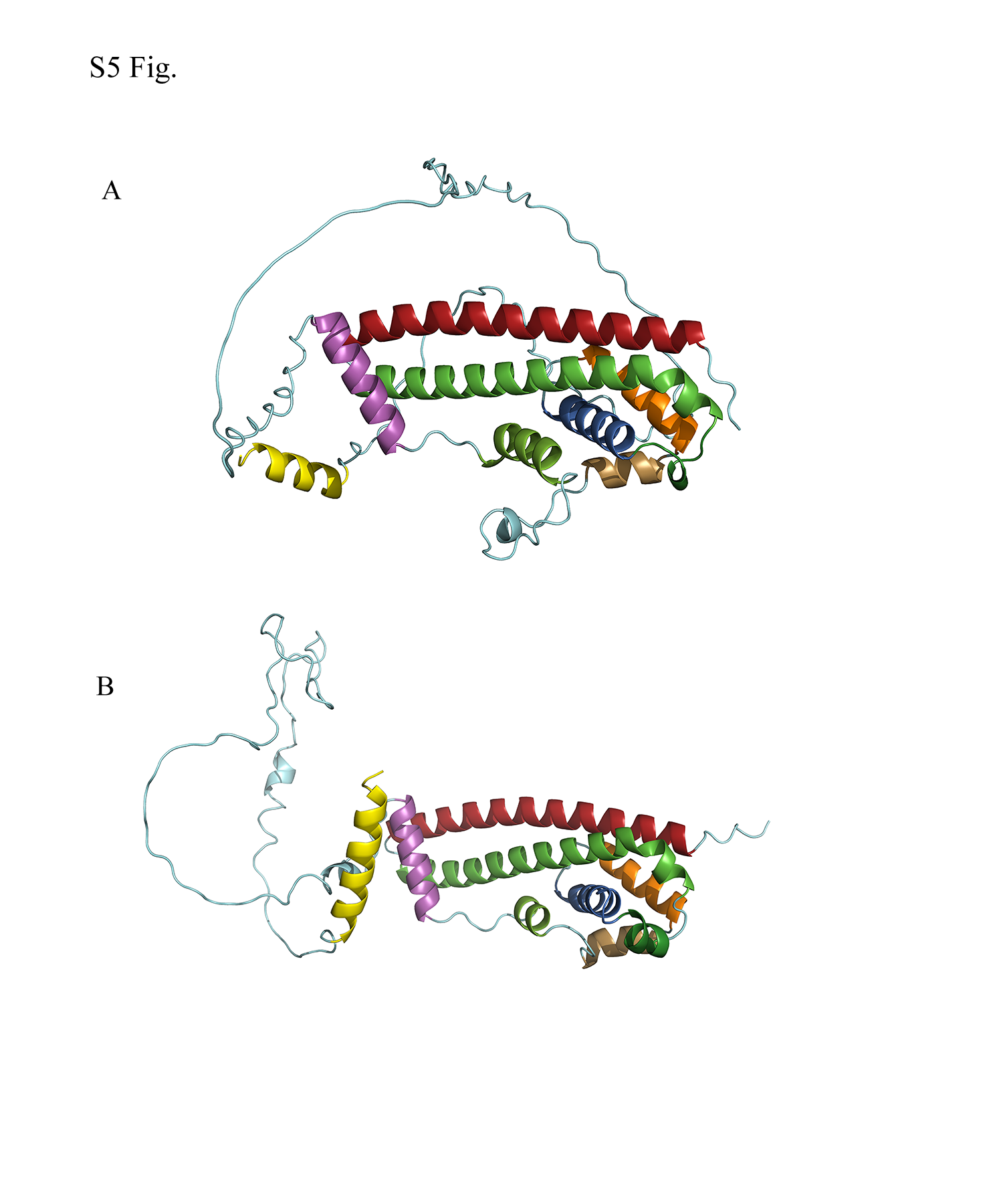

### Supplementary Figure 6

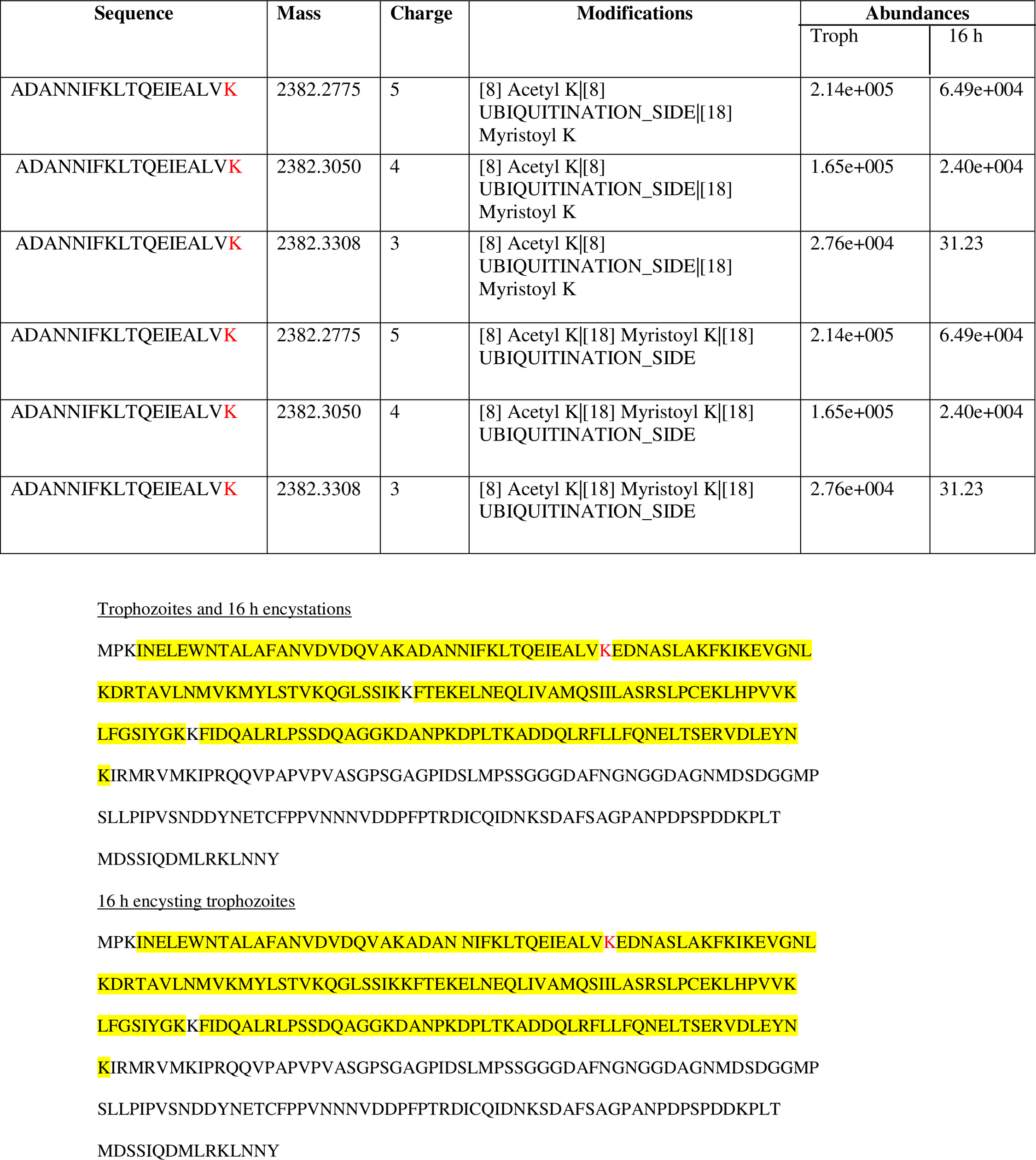
